# Markovian State Models uncover Casein Kinase 1 dynamics that govern circadian period

**DOI:** 10.1101/2025.01.17.633651

**Authors:** Clarisse Gravina Ricci, Jonathan M. Philpott, Megan R. Torgrimson, Alfred M. Freeberg, Rajesh Narasimamurthy, Emilia Pécora de Barros, Rommie Amaro, David M. Virshup, J. Andrew McCammon, Carrie L. Partch

**Author notes:** Corresponding authors (C.G.R.), (C.L.P.).

## Abstract

Circadian rhythms in mammals are tightly regulated through phosphorylation of Period (PER) proteins by Casein Kinase 1 (CK1, subtypes δ and ε). CK1 acts on at least two different regions of PER with opposing effects: phosphorylation of phosphodegron (pD) regions leads to PER degradation, while phosphorylation of the Familial Advanced Sleep Phase (FASP) region leads to PER stabilization. To investigate how substrate selectivity is encoded by the conformational dynamics of CK1, we performed a large set of independent molecular dynamics (MD) simulations of wildtype CK1 and the *tau* mutant (R178C) that biases kinase activity toward a pD. We used Markovian State Models (MSMs) to integrate the simulations into a single model of the conformational landscape of CK1 and used Gaussian accelerated molecular dynamics (GaMD) to build the first molecular model of CK1 and the unphosphorylated FASP motif. Together, these findings provide a mechanistic view of CK1, establishing how the activation loop acts as a key molecular switch to control substrate selectivity. We show that the *tau* mutant favors an alternative conformation of the activation loop and significantly accelerates the dynamics of CK1. This reshapes the binding cleft in a way that impairs FASP binding and would ultimately lead to PER destabilization and shorter circadian periods. Finally, we identified an allosteric pocket that could be targeted to bias this molecular switch. Our integrated approach offers a detailed model of CK1’s conformational landscape and its relevance to normal, mutant, and druggable circadian timekeeping.

**Statement of Significance:** Disruption of circadian rhythms alters the temporal orchestration of vital cellular processes and increases the propensity for sleep disorders, metabolic disease, and cancer. Circadian rhythms are generated by a vast gene expression program controlled at the cellular level by a molecular clock comprised of dedicated clock proteins. Amongst the essential protein characters is Casein kinase 1 (CK1), which acts on multiple clock protein substrates. A delicate balance of CK1 activity on these substrates is crucial for proper circadian timekeeping, highlighting CK1 as a promising drug target to tune clock timing. This work aims to identify the conformational landscape of CK1 that underlies its substrate specificity and provide molecular insight for pharmacologic development that could modulate CK1 function for those suffering from clock-related syndromes.

## Introduction

One of the greatest achievements of life on Earth is the ability of organisms to anticipate the terrestrial cycles of light and darkness. From prokaryotes to mammals, life forms take advantage of day and night to perform cellular tasks in a coherent way by following an internal clock [1–5]. While the biological clock synchronizes to the solar day by making use of external cues, its intrinsic ‘ticking’ pace is dictated by an internal core oscillator present in nearly every cell and displaying a period of ∼24 hours [6]. In mammals, the core oscillator consists of an interlocked transcription/translation feedback loop that generates daily oscillations in gene expression [3, 7]. This results in a circadian (about a day) expression of proteins involved in behavior, development, metabolism, DNA repair, and more [6]. Mutations causing the intrinsic period to be significantly different from ∼24 hours can prevent organisms from successfully synchronizing their clocks to the solar day. This results in social jetlag and sleep disorders, which in the long run, can interfere with our immune response [8, 9] and trigger pathologies such as metabolic syndrome, diabetes, and cancer [6, 10–18]. In this scenario, understanding the molecular underpinnings of the clock could unlock new pharmacological targets to treat a wide range of diseases.

In humans, the transcription/translation feedback loop is formed by a transcription activator complex, CLOCK:BMAL1, and a repressor complex formed by Period proteins (PER1 and PER2), Cryptochromes (CRY1 and CRY2), and the protein Casein Kinase 1 (subtypes δ and ε, hereafter jointly referred to as CK1) (Figure 1A) [19–23]. The CLOCK:BMAL1 dimer activates the transcription of many circadian-controlled genes, including those of their repressors (PERs and CRYs), leading to daily oscillations in repressor expression. Because PER is the stoichiometric limiting factor in the assembly of the CK1:PER:CRY repressor complex [24], its abundance and stability are correlated with the duration of the intrinsic circadian period. The molecular mechanisms regulating the life span of PER proteins thus provide a direct link to circadian period. At the very center of this switch is CK1, which modulates PER stability through post-translational modifications [25–30] and is thought to confer temperature insensitivity to circadian rhythms [31, 32].

**Figure 1.**
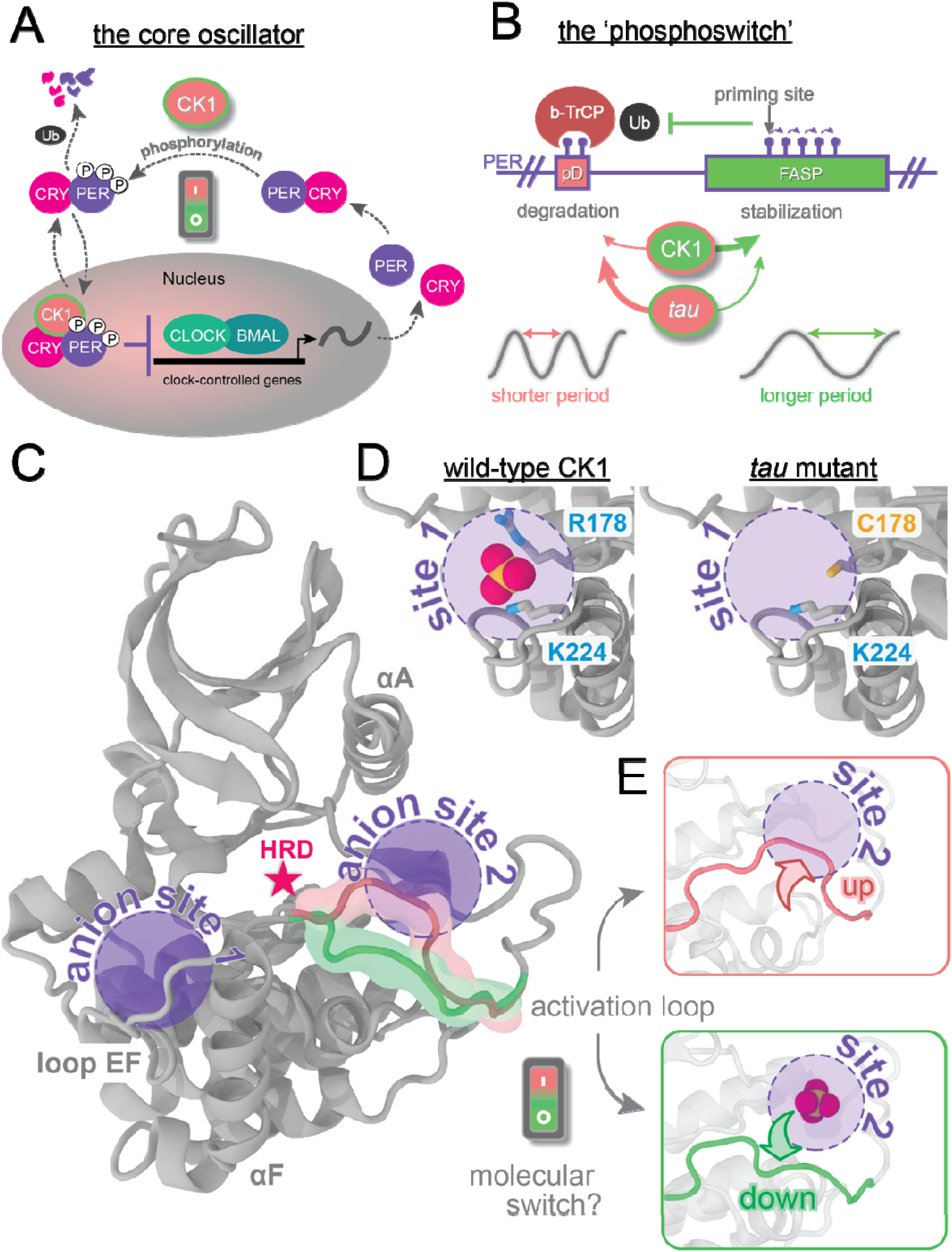
The CK1 activation loop acts as a molecular switch to control PER phosphorylation and circadian period. A) Simple schematic view of the transcriptional-translational feedback loop at the core of the human clock. B) The circadian period is finely regulated by a phosphoswitch mechanism controlling PER stability. C) Apo structure of CK1δ (PDB 1CKJ), highlighting: the location of the catalytic HRD motif (red star); the two conserved anion binding sites (purple circles), and the alternative conformations of the activation loop (pink and green). D) Sulfate anions bind to the first anion site via two positively charged clamps (R178 and K224). The R178C mutation in *tau* impairs the ability of this site to bind anions. E) Alternative conformations of the activation loop in *tau* (PDB 6PXN), showing that the loop up conformation (pink) sterically blocks the second anion site.

CK1 phosphorylates at least two different regions on PER with antagonist effects on its stability in a phosphoswitch mechanism (Figure 1B) [33–35]. Phosphorylation at phosphodegron (pD) regions recruits the E3 ubiquitin ligase, β-TrCP, and leads to PER degradation, with the ultimate effect of shortening the circadian period [36–40]. On the other hand, phosphorylation of five consecutive serines in the FASP region [25] found within the CK1-binding domain of PER leads to its stabilization through feedback inhibition that attenuates kinase activity at the pD [41]. Mutation of the first serine in this region causes abnormally short circadian periods of ∼20 hours and Familial Advanced Sleep Phase [36, 42]. Other mutations in CK1/CK1-orthologs [43–51] or in its phosphorylation sites on PER/PER-orthologs [34, 45] can induce remarkable changes to the intrinsic circadian period, underscoring the central importance of the phosphoswitch for clock timing. Not surprisingly, CK1 is gaining traction as an effective pharmacological target for dysfunctional clocks [27, 52]. ATP-competitive CK1 inhibitors capable of lengthening the circadian rhythm have been identified [27, 53–60], but, unlike many mutations in the enzyme, no small molecules are capable of shortening the period [44, 47, 49, 50]. To develop highly specific circadian drugs that diversify our ability to modulate period length, more information is needed on the molecular factors dictating CK1 regulation and substrate selectivity.

CK1 is a ubiquitously expressed serine/threonine kinase with activity against a broad variety of substrates in all cell types [61, 62]. Orthologs of CK1 have been implicated in timekeeping in a variety of eukaryotic organisms ranging from green algae to humans [47, 49, 63, 64], denoting a well-conserved and far-reaching role across species. CK1 displays the typical two-lobed kinase architecture; however, it is not regulated by phosphorylation of its activation loop (Figure 1C)[61, 65, 66]. Instead, the kinase domain of CK1 is constitutively active [62, 67, 68] and two highly conserved anion binding pockets play important regulatory roles, including substrate selectivity and recognition [43, 66, 69]. CK1 preferentially acts on negatively charged or primed substrates, such as those with a D/E/pSxxS consensus motif [66, 70]. Interestingly, the CK1-dependent phosphorylation sites that form the phosphoswitch in PER––pD and FASP––do not display this consensus motif and are slow, rate-limiting steps for the circadian clock [25, 33]. Thus far, little is known about the conformational mechanism by which CK1 balances its activity between the pD and FASP regions.

The CK1-based circadian mutant *tau* [71] shortens the circadian period by approximately 4 hours [48, 51]. In this mutant, an arginine in the first anion binding pocket is replaced by a cysteine (R178C) (Figure 1D), impairing the pocket’s ability to bind negatively charged groups in primed or acidic substrates [69, 71]. This site lies near loop EF (L-EF), a flexible region generally implicated in substrate binding by kinases [66, 70] and, in the case of CK1, specifically required for temperature-compensation [31]. We showed that *tau* simultaneously reduces the activity on the FASP region and increases the activity on the pD site, inverting substrate selectivity on PER relative to the wild-type enzyme (Figure 1B) [71]. This makes the *tau* mutant an ideal system to study the conformational mechanisms of how CK1 toggles the phosphoswitch. Supported by molecular dynamics (MD) simulations, the crystal structure of *tau* revealed two alternative conformations of the activation loop (‘up’ and ‘down’, Figure 1E), hinting at a two-state conformational switch at the core of CK1 substrate selectivity. Interestingly, the less frequent ‘loop up’ conformation of the wild-type CK1 sterically abolishes the second anion binding pocket, which could be a determining factor for substrate binding and/or product inhibition [71, 72].

To investigate how substrate selectivity is encoded in the conformational dynamics of CK1, here we performed a large set of unbiased MD simulations of wild-type (WT) CK1 and the *tau* mutant. These simulations were integrated into a Markov State Model (MSM) [73–76] to describe the free energy landscape bridging ‘loop up’ and ‘loop down’ conformations in CK1. MSMs have been emerging as a powerful framework to uncover slow dynamics in complex biomolecular systems [77–79] and to quantify differences between protein ensembles [80–82], many times with significant implications for drug discovery and design [83–87]. Here, we additionally used Gaussian accelerated molecular dynamics (GaMD) [88–91] to boost the sampling of the CK1 conformational landscape and to build the first molecular model of the interaction of CK1 with the priming motif of FASP. Altogether, we find that the *tau* mutant accelerates CK1 dynamics, favors the ‘loop up’ conformation of the activation loop, and reshapes the binding cleft in a way that hinders FASP binding and ultimately destabilizes PER. Our combined approach provides a comprehensive model of the conformational landscape of CK1 and its implications for circadian timekeeping by establishing the activation loop as a key molecular switch for substrate selectivity and identifying a potential allosteric pocket that could be targeted to shorten the circadian period.

## Materials and Methods

### Molecular dynamics simulations

All information about the set-up, execution and analysis of the molecular dynamics simulations is available in the Supporting Materials online.

### Expression and purification of recombinant proteins

All plasmid purification was carried out in *Escherichia coli* DH5a cells. Proteins were expressed from a pET22-based vector in *Escherichia coli* BL21 (DE3) Rosetta2 cells (Sigma Aldrich) based on the Parallel vector series [92]. All FASP peptides were expressed downstream of an N-terminal TEV-cleavable His-NusA tag. Human CK1δ catalytic domains (CK1, residues 1–317) were all expressed in BL21 (DE3) Rosetta2 cells (Sigma Aldrich) with a TEV-cleavable His-GST tag. Mutations were made using standard site-directed mutagenesis protocols and validated by sequencing. All proteins and peptides expressed from Parallel vectors have an N-terminal vector artifact (GAMDPEF) remaining after TEV cleavage and the peptides have a tryptophan and polybasic motif (WRKKK) following the vector artifact. Cells were grown in LB media (for natural abundance growths) or M9 minimal media with the appropriate stable isotopes (i.e., ^15^N/^13^C for NMR) as done before [25] at 37°C until the O.D._600_ reached ∼0.8; expression was induced with 0.5 mM IPTG, and cultures were grown for approximately 16–20 hours more at 18°C.

For CK1 protein preps, cells were lysed in 50 mM Tris pH 7.5, 300 mM NaCl, 1 mM TCEP, and 5% glycerol using a high-pressure extruder (Avestin) or sonicator (Fisher Scientific) on ice. HisGST-CK1 fusion proteins were purified using Glutathione Sepharose 4B resin (GE Healthcare) using standard approaches and eluted from the resin using Phosphate Buffered Saline with 25 mM reduced glutathione. His-TEV protease was added to cleave the His-GST tag from CK1 at 4°C overnight. Cleaved CK1 was further purified away from His-GST and His-TEV using Ni-NTA resin (Qiagen) and subsequent size exclusion chromatography on a HiLoad 16/600 Superdex 75 prep grade column (GE Healthcare) in 50 mM Tris pH 7.5, 200 mM NaCl, 5 mM BME, 1 mM EDTA, and 0.05% Tween 20. Purified CK1 proteins used for *in vitro* kinase assays were run on size exclusion columns or buffer exchanged into storage buffer (50 mM Tris pH 7.5, 100 mM NaCl, 1 mM TCEP, 1 mM EDTA, and 10% glycerol) using an Amicron Ultra centrifugal filter (Millipore) and frozen as small aliquots in liquid nitrogen for storage at −80°C.

For human PER2 FASP peptide preps, cells were lysed in a buffer containing 50 mM Tris pH 7.5, 500 mM NaCl, 2 mM TCEP, 5% glycerol and 25 mM imidazole using a high-pressure extruder (Avestin) or sonicator on ice (Fisher Scientific). His-NusA-FASP fusion proteins were purified using Ni-NTA resin using standard approaches and eluted from the resin using 50 mM Tris pH 7.5, 500 mM NaCl, 2 mM TCEP, 5% glycerol and 250 mM imidazole. His-TEV protease was added to cleave the His-NusA tag from the PER2 peptides at 4°C overnight. The cleavage reaction was subsequently concentrated and desalted into low imidazole lysis buffer using a HiPrep 26/10 Desalting column. Peptides were purified away from His-NusA and His-TEV using Ni-NTA resin with 50 mM Tris pH 7.5, 500 mM NaCl, 2 mM TCEP, 5% glycerol and 25 mM imidazole. Peptides were purified by size exclusion chromatography on a HiLoad 16/600 Superdex 75 prep grade column, using NMR buffer (25 mM MES pH 6.0, 50 mM NaCl, 2 mM TCEP, 10 mM MgCl_2_) or 1x kinase buffer (25 mM Tris pH 7.5, 100 mM NaCl, 10 mM MgCl_2_, and 2 mM TCEP) for NMR or ADP-Glo kinase assays, respectively.

### NMR kinase assays

NMR spectra were collected on a Varian INOVA 600 MHz or a Bruker 800 MHz spectrometer equipped with a ^1^H, ^13^C, ^15^N triple resonance z-axis pulsed-field-gradient cryoprobe. Spectra were processed using NMRPipe [93] and analyzed using CCPNmr Analysis [94]. Backbone resonance assignments were determined previously [95]. NMR kinase reactions were performed at 25°C with 150 µM ^15^N-human PER2 FASP, 2.5 mM ATP and 2 µM CK1. SOFAST HMQC spectra (data acquisition = 5 min) were collected at the indicated timepoints, or HSQC spectra were collected on quenched samples (after addition of EDTA to final concentration of 20 mM) at the indicated timepoints, and the relative peak volumes were calculated and normalized as described previously [25].

### ADP-Glo kinase assays

Kinase reactions were performed on the indicated recombinant peptides (FASP WT or alanine mutants) using the ADP-Glo kinase assay kit (Promega) according to manufacturer’s instructions. All reactions were performed in 30 μL volumes using 1x kinase buffer (25 mM Tris pH 7.5, 100 mM NaCl, 10 mM MgCl_2_, and 2 mM TCEP) supplemented with ATP and substrate peptides. To determine apparent kinetic parameters (K_M_), duplicate reactions with 100 µM ATP and 200 nM CK1 kinase were incubated in 1xkinase buffer at room temperature for 1 hour with the indicated amount of substrate peptide (and repeated for n = 2 independent assays). 5 µL aliquots were taken and quenched with ADP-Glo reagent after the 1 hour incubation, and Luminescence measurements were taken at room temperature with a SYNERGY2 microplate reader (BioTek) in 384-well microplates. Data analysis was performed using Excel (Microsoft) or Prism (GraphPad).

### Radioactive kinase assays

PER FASP region peptides were synthesized and purified to 95% or higher (SABio). Two independent reaction mixtures of 50 μL containing 200 μM of the FASP or in reaction buffer (25 mM Tris pH 7.5, 7.5 mM MgCl_2_, 1 mM DTT, 0.1 mg/mL BSA) were preincubated for 5 minutes with or without 20 nM CK1 (for primed FASP) or 200 nM CK1 (for unprimed FASP) and the reaction was started by addition of 750 μM of UltraPure ATP (Promega) containing 1-2 μCi of ⍰-^32^ p ATP (Perkin Elmer). After incubation of the reaction mix at 30 °C, an 8 μL aliquot of the reaction mix was transferred to P81 phosphocellulose paper (Reaction Biology Corp) at the indicated timepoints. The P81 paper was washed three times with 75 mM of orthophosphoric acid and once with acetone. The air-dried P81 paper was counted for P_i_ incorporation using a scintillation counter (Perkin Elmer) by Cherenkov counting. Results shown are from four independent assays.

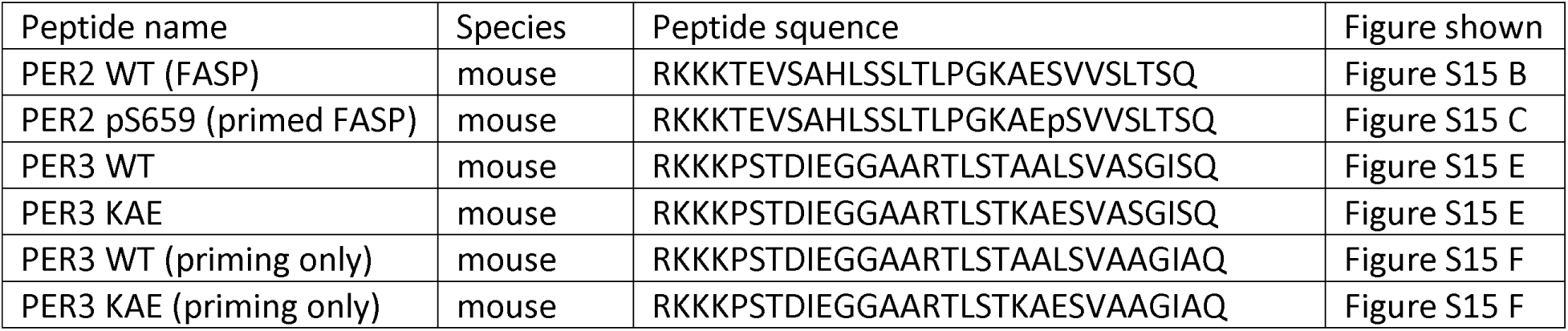

## Results

### Mapping the conformational landscape of CK1

#### The activation loop and loop EF are the slowest loops in CK1

To characterize the dynamics of WT CK1 and the *tau* mutant, we ran multiple all-atom molecular dynamics simulations totaling 21 μs for each protein system (Figure S1). We then employed the MSM framework to integrate these trajectories into a single model describing the conformational free energy landscape of CK1 in wild-type and *tau* Markov State Models can derive long timescale dynamics from a large number of relatively short MD simulations by (i) dividing the conformational space into a large number of discrete microstates, (ii) using the MD data to count transitions between states after a specified lag time; and finally iii) estimating a transition matrix that describes the dynamics of the system in the discretized conformational space [73, 96]. An important step in model building thus consists in selecting relevant features to discretize the conformational space. For our model, we started by selecting pair-wise distances involving functionally important regions of CK1 and applied time-lagged independent component analysis (tICA) [97] to reduce these features to a smaller number of collective variables representing the slowest modes of motions (TICs). We found that the slowest and dominant mode of motion in CK1 involves the activation loop (TIC 1) and that interconversion between the ‘loop up’ and ‘loop down’ conformations happens at long scales, elucidating an important regulatory role for this loop (Figure S2). The second slowest motion (TIC 2) involves L-EF, near the first anion site, which has been previously implicated in substrate binding and temperature compensation mechanisms [31, 66, 70]. Based on these results, we selected five pair-wise distances that were jointly combined by tICA to create the final MSMs. Further methodological details on construction of the models are provided in Methods and Supplementary Information (Figures S3-S5).

#### Tau accelerates and inverts the conformational equilibrium involving the activation loop

MSM-based conformational landscapes for WT CK1 and the *tau* mutant reveal that each system displays three preferred states (Figure 2A). Two of these states are roughly equivalent in WT and *tau* (states I/I’ and III/III’), while states II and IV’ are exclusive of WT and *tau* respectively. Visual inspection supported by pair-wise distances and RMSD (Figure S6) reveal that the states differ mainly with respect to the conformation of the activation loop (up, down, or intermediate) and the conformational state of L-EF (folded or unfolded) (Figure 2B). States I/I’ and III/III’ correspond to ‘loop up’ and ‘loop down’ conformations of the activation loop, respectively, in relatively good agreement with corresponding crystallographic structures. The remaining states display the L-EF in an unfolded state, with the activation loop either adopting an intermediate conformation (state II, in WT) or the ‘loop down’ conformation (state IV’, in *tau*).

**Figure 2.**
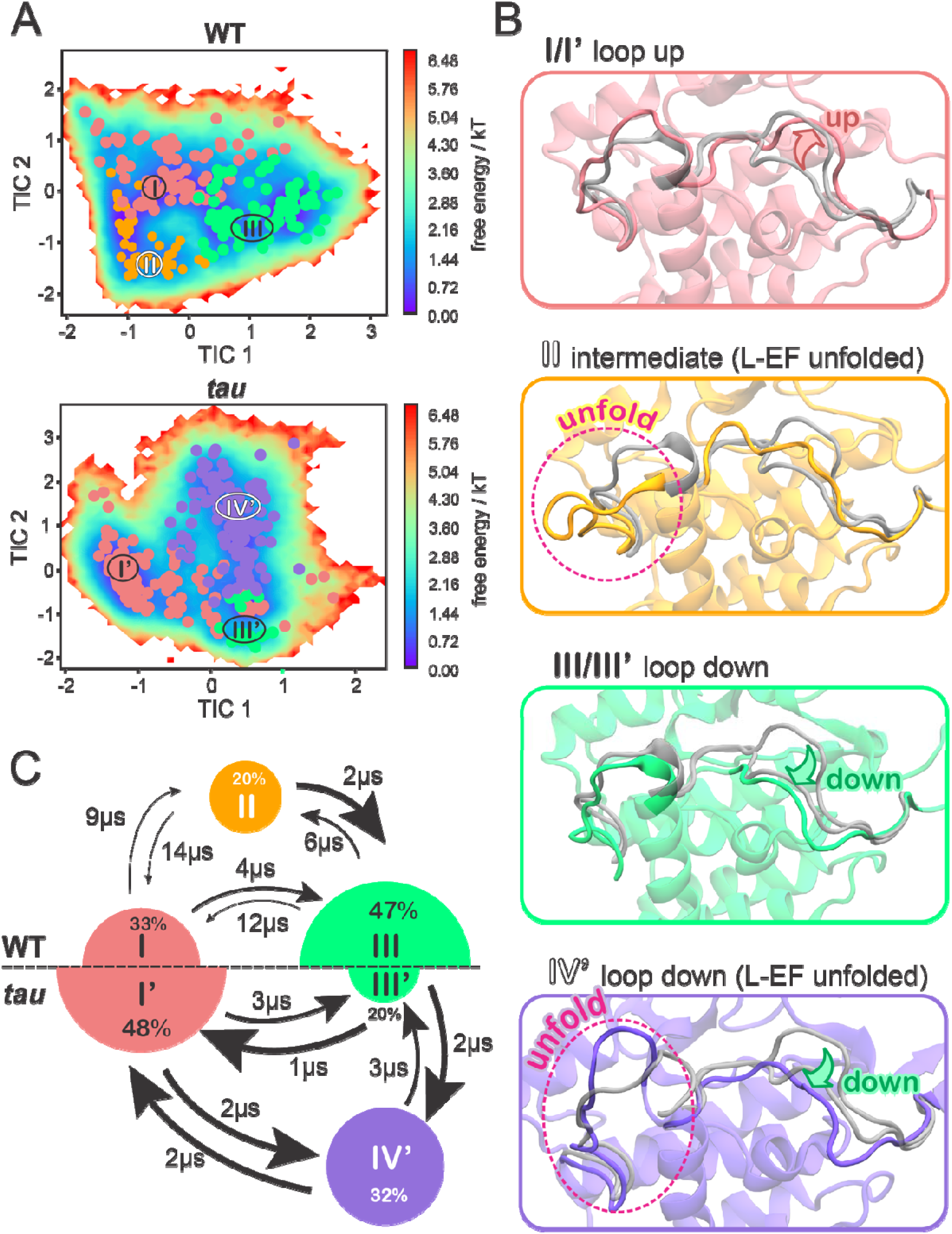
The equilibrium between three preferred conformational states is accelerated and inverted by the CK1 *tau* mutant. A) Free energy landscapes of the WT CK1 (top) and *tau* mutant (bottom) in terms of the slowest tICA components, with microstates clustered into meta-stable states (identified by roman numerals). B) Representative conformations of each meta-stable state. For comparison, x-ray conformations of the activation loop and L-EF are superimposed (transparent gray; loop down PDB 1CKJ, loop up PDB 6PXN). C) Equilibrium populations of meta-stable states and MFPTs between states for the WT (top) and *tau* (bottom) protein systems. Radius of the circles is proportional to the equilibrium population percentage and thickness of the arrows is proportional to transition rates between states. The numbers next to the arrows indicate MFPTs.

Populations derived from the MSM uncover a slow equilibrium between ‘loop up’ (state I) and ‘loop down’ (state III) conformations in the wild-type kinase, with a clear preference for the ‘loop down’ conformation (Figure 2C, top). The observation of an additional metastable state (II) in which the activation loop adopts a ‘halfway’ conformation suggests the existence of an intermediate state connecting the two main states (I and III) in the wild-type enzyme. This intermediate state also suggests that flipping of the activation loop requires disorganization of L-EF, which displays significantly more conformational freedom in state II (Figure S6).

Interestingly, the *tau* mutation inverts the conformational equilibrium characteristic of WT, stabilizing the activation loop in the ‘up’ conformation (state I’) and destabilizing the ‘down’ conformation (Figure 2C, bottom), which is almost always accompanied by unfolding of L-EF (state IV’). In addition, flipping of the activation loop between up and down conformations does not involve a long-lived intermediate or ‘midway’ conformation. Such inversion of the conformational landscape strongly supports the activation loop as a key molecular switch controlling the circadian period since the *tau* mutant is known for inverting CK1 selectivity for its circadian substrates. Our model also recapitulates previous findings that the *tau* mutation destabilizes the first anion site, facilitating the unfolding of L-EF, but *only* when the activation loop is down (state IV’) [71]. When the activation loop is up, L-EF remains folded and displays WT-like dynamics (state I’), supporting that the activation loop exerts allosteric control over the dynamics of L-EF.

Apart from equilibrium populations, MSMs also provide transition rates between states, informing on the kinetics of the system. By comparing mean first passage times (MFPT) between states (Figure 2C, arrows), we find that the *tau* mutation significantly accelerates the conformational transitions between states. Interconversion between loop up and down conformations in the *tau* mutant (I’ ⍰ IV’ ≈ 2 μs) is at least two times faster than in the wild-type enzyme (I ⍰ III ≈ 4-12 μs), indicating that the *tau* mutation significantly reduces the energy barriers associated with flipping of the activation loop. This agrees with the lack of intermediate ‘midway’ conformations in the *tau* mutant. *Tau* also accelerates unfolding of L-EF, which in state IV’, adopts a wide range of conformations ranging from collapsed to fully extended (see Figure S6B).

#### The role of Gly^175^ in the conformational dynamics of the activation loop

The activation loop in CK1 is preceded by a conserved glycine at position 175. In other serine/threonine kinases, a backbone flip of a glycine at this conserved position has been linked to conformational changes of the activation loop [98]. In CK1, x-ray structures suggested that Gly^175^ backbone could work as a ‘hinge’ controlling the conformation of the activation loop (Figure 3A) [66, 71]. To investigate this hypothesis, we built MSMs based solely on the backbone angles of Gly^175^ (Figures S7-8).

**Figure 3.**
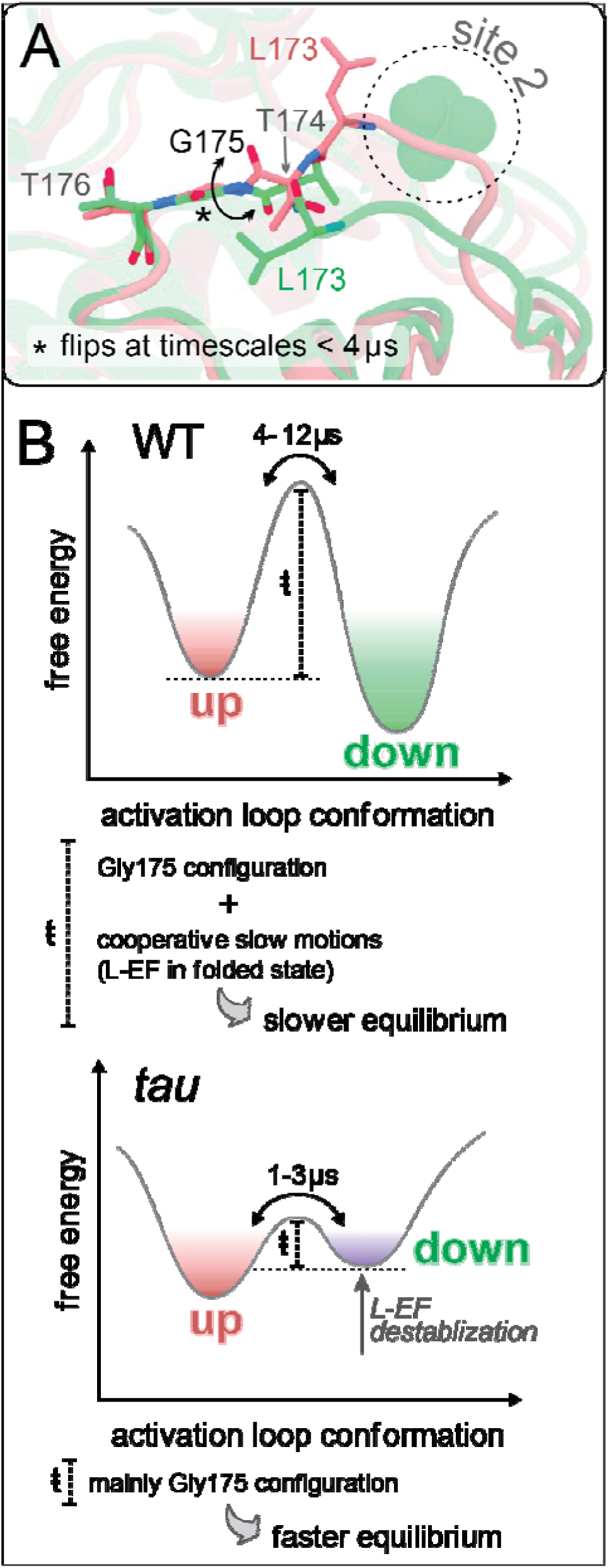
The role of Gly^175^ for the conformational dynamics of the activation loop. A) View of x-ray structures highlighting the different configurations adopted by Gly^175^ in ‘loop up’ and ‘loop down’ conformations of the CK1 activation loop (PDB 6PXN and 1CKJ, respectively). B) Schematic representation of the conformational landscape involving the activation loop, based on MSM-derived kinetics and stability of the two most populated states in each system (WT CK1, top; and *tau*, bottom).

The resulting Gly^175^-based MSMs revealed no significant correlation between the configuration of Gly^175^ and the conformation of the activation loop in WT CK1 (Figure S9). Thus, while the configuration of Gly^175^ might determine the most energetically stable conformation of the activation loop in low entropy scenarios (crystalline state), its importance appears to be overcome by other factors when the protein is in solution. Surprisingly, the *tau* mutant displays a moderate correlation between φ^Gly175^ and the activation loop (φ^Gly175^ < 0° favors ‘loop up’ and φ^Gly175^ > 0° favors ‘loop down’), indicating that Gly^175^ backbone is more determinant of the conformation of the activation loop in the *tau* mutant than in the wild-type enzyme.

In part, this can be explained by the energetic barriers separating states in the conformational landscape of CK1. In the wild-type enzyme, a configurational torsion of Gly^175^ alone is not enough to overcome the high energy barriers associated with the flip of the activation loop. In the *tau* mutant, these barriers are decreased by disruption of the first anion binding site and unfolding of L-EF, allowing the conformational dynamics of the activation loop to be influenced more heavily by the backbone configuration of Gly^175^ This is in excellent agreement with MFPTs provided by the MSMs which show that torsional transitions of Gly^175^ happen at timescales faster than 4 μs (Figure S8A), the same time-scale range at which the activation loop flips in the *tau* mutant (see Figure 2C). In WT CK1, loop transitions happen at much longer timescales (4-12 μs) (see Figure 2C), indicating that the conformational landscape in the wild-type enzyme is governed by slow cooperative motions, likely related to the conformational state of L-EF (Figure 3B).

### A molecular model of FASP interactions involved in priming

To gain a better understanding on how the conformational landscape of CK1 is linked to its activity on circadian substrates, we decided to model the interaction between CK1 and the FASP region of PER2 (Figure 4A). We based our initial model on a recently published crystallographic structure of CK1 bound to TAp63α (Figure S10) [72]. As with FASP, TAp63α also undergoes sequential phosphorylation by CK1, with the difference that the priming is achieved by another kinase, CDK2 [99]. We refined our model with a combined set of 2 μs of accumulated GaMD simulations followed by additional 2 μs of accumulated cMD simulations (Figure S11, details in SI).

**Figure 4.**
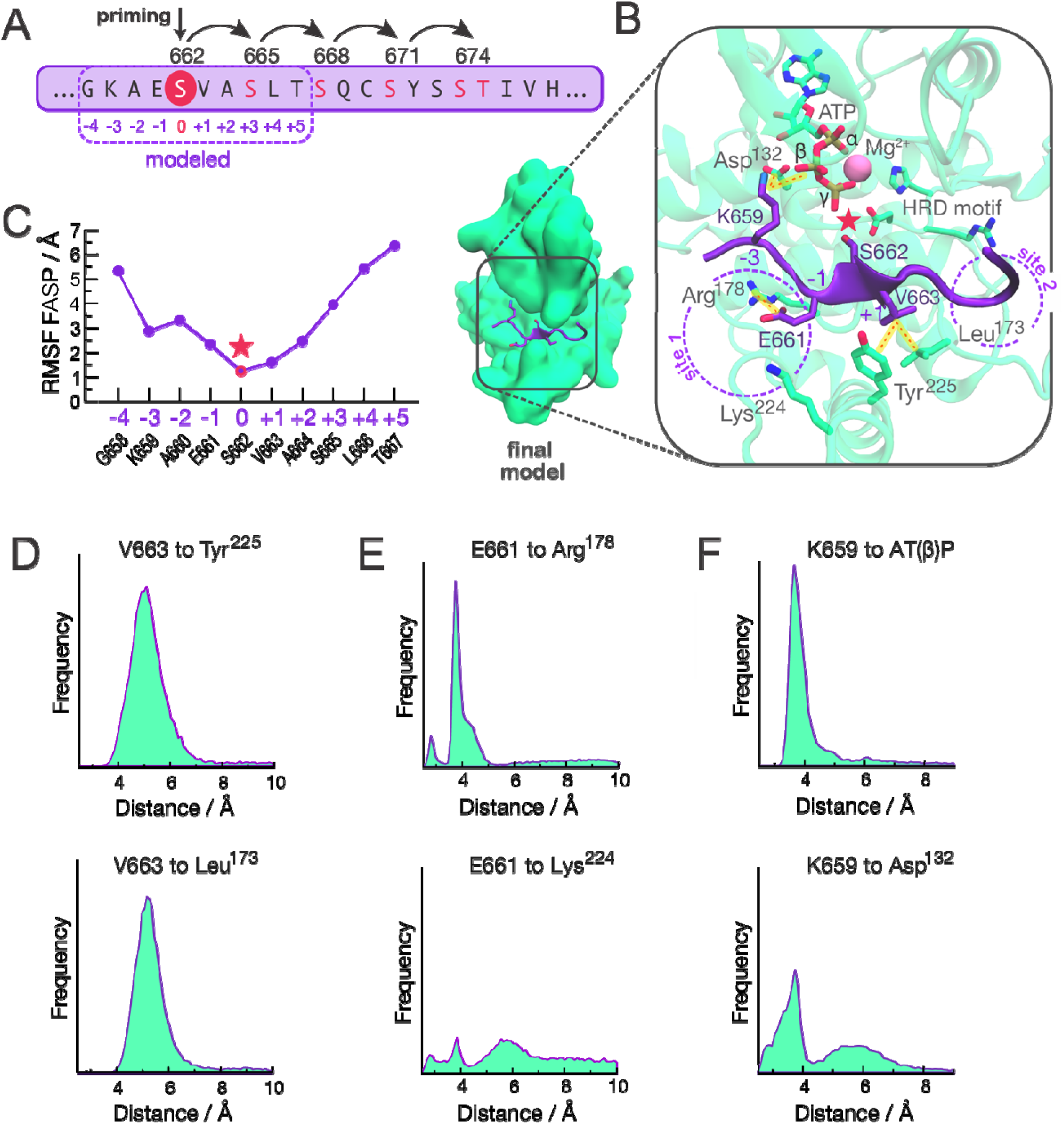
Molecular model of the interaction between CK1 and unphosphorylated FASP. A) The FASP region of human PER2. Residue numbers with arched arrows represent serines that are sequentially phosphorylated by CK1. Positions listed below (−4 - +5) are relative to priming serine, S662. B) Representative structure of the final model with the unphosphorylated FASP peptide (purple) and CK1 (green). For clarity, we use one-letter amino acid code for residues of the substrate (FASP) and three-letter code for residues of the kinase CK1. Pink star represents the priming event of S662. C) Atomic fluctuations of the bound FASP based on accumulated molecular dynamics trajectories. D-F) Distance-based interaction histograms involving V663 at position +1 (panel D), E661 at position −1 (panel E), and K659 at position −3 (panel F) with CK1 residues or ATP.

#### FASP binds to the loop down conformation

Our final model consists of a FASP-bound conformation in which the priming serine (S662 in human PER2) is well positioned to accept the γ phosphate from ATP, with the activation loop in the ‘down’ conformation (Figure 4B and Figure S11B). V663 (at position +1) appears to play a key role in anchoring the backbone of FASP into the active site (see Figure 4B), displaying low atomic fluctuations (Figure 4C) and engaging in persistent hydrophobic interactions with Tyr^225^ in helix αF and with Leu^173^ in the activation loop (Figures 4D).

#### Electrostatic interactions upstream of the priming site of FASP contribute to substrate binding

While residues downstream of the priming site display high mobility freedom and a lack of persistent interactions with CK1, the upstream region of FASP is less dynamic and likely to contribute to substrate binding (Figure 4C). We noticed that the first anion binding site is partially occupied by E661 (at position-1) which displays persistent electrostatic interactions with Arg^178^ but not as much with Lys^224^ (Figure 4E). In the *tau* mutant, Arg^178^ is replaced with a cysteine, likely disrupting this interaction. We also found that K659 (at position −3) invariably forms a salt bridge with the β phosphate of ATP, oftentimes assisted by an additional salt bridge with an Asp^132^ (just downstream of the catalytic HRD motif) (Figure 4F).

### Biochemical validation of the PER FASP priming model

To support our model of unprimed FASP bound to CK1, we performed a series of biochemical experiments to assess the relative contribution of residues near the FASP priming site as molecular determinants of CK1 priming activity. As we have done previously [25, 41, 71], we used an NMR-based kinase assay to measure priming activity within human PER2 FASP peptides with site-specific resolution. In agreement with our model, introducing alanine mutations to the −3, −1, or +1 positions of FASP led to a decrease in priming activity (Figure 5A, Figure S14).

**Figure 5.**
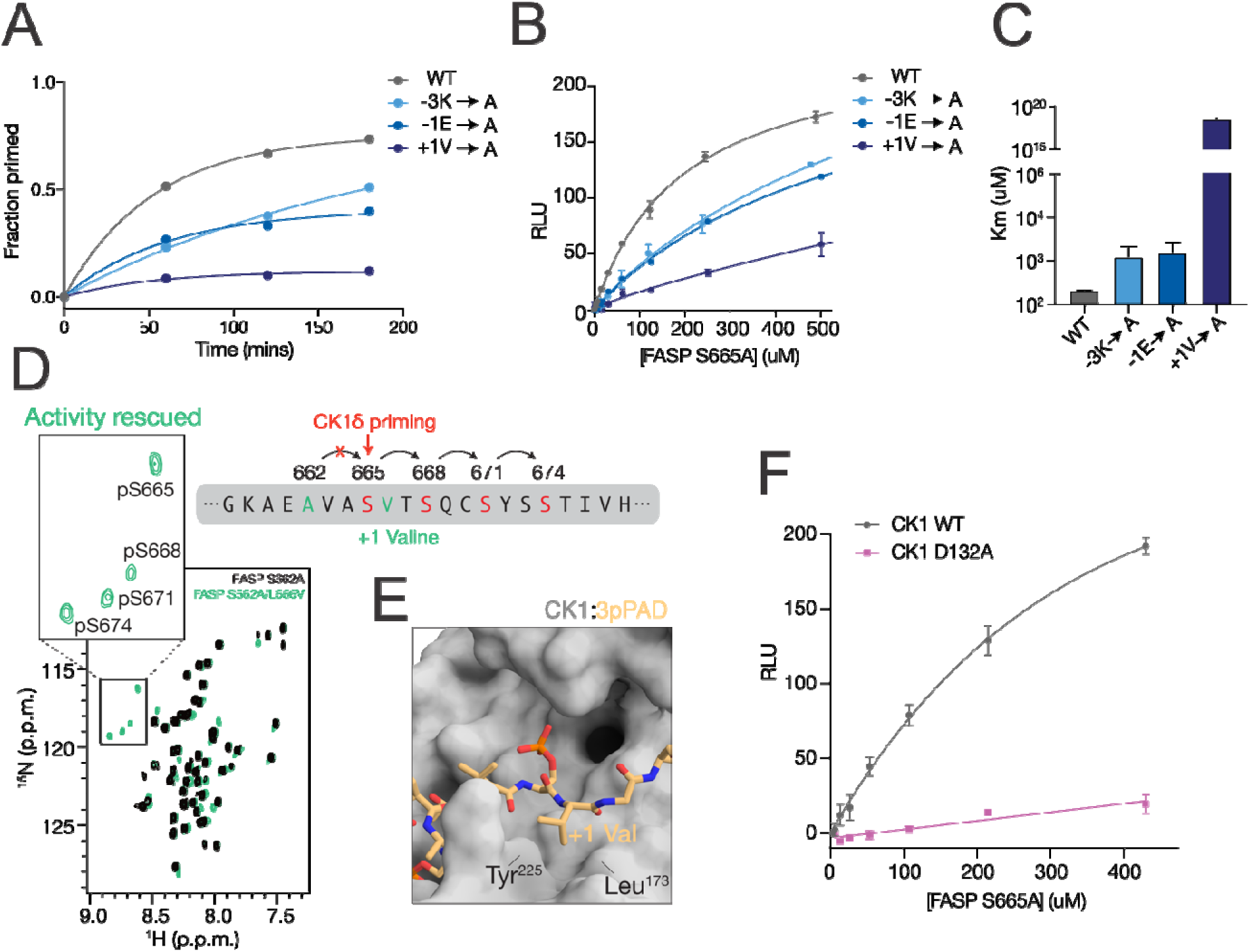
Biochemical validation of the model for non-consensus FASP priming. A) NMR-based timecourse kinase assay quantifying the increase in peak volume corresponding to the phosphopeak of the priming serine in human PER2 FASP (S662), taken from a series of ^15^N-^1^H HSQC spectra (Figure S14). B) ADP-Glo titration assay comparing WT FASP and alanine mutants at positions −3, −1, and +1 in the ‘priming only’ background (S665A mutation to halt sequential kinase activity, see Figure S15A). C) Quantification of the mean and SD of the KM from replicate titration experiments as shown in panel B, n=2. D) ^15^N-^1^H HSQC spectra comparing a ‘priming disrupted’ FASP peptide (S662A, black) and a ‘priming rescued’ FASP peptide (S662A/L666V, teal) in the presence of CK1. Zoom shows the region of the spectra where phosphoserines appear. E) Zoom of co-crystal structure o CK1 and a triply phosphorylated Tap63α PAD peptide, 3pPAD, highlighting the active site and +1 hydrophobic pocket region (PDB 6RU8). The +1 valine residue is inserted between the small hydrophobic pocket created between Tyr^225^ and Leu^173^ of CK1 when the activation loop is in the downward conformation. F) ADP-Glo titration of the ‘priming only’ (S665A) FASP peptide comparing CK1 WT (black) and D132A (purple).

To get a further sense of how these mutations might contribute to the binding of unprimed FASP, we introduced an alanine mutation at the +4 Ser (S665) to limit kinase activity to just the priming site (Figure S15A). We then performed substrate titrations of FASP peptides with alanine mutations at the −3, −1, and +1 positions in this ‘priming only’ (S665A) background by ADP-Glo assay. Similar to the NMR assay, these mutations led to an increase in the Michaelis-Menten constant (K_M_) suggesting reduced affinity of the unprimed FASP substrate for the kinase (Figures 5B-C). Mutation of the +1 valine in FASP had the largest decrease in kinase activity, in agreement with lower overall atomic fluctuations displayed by this residue in our MD model (Figure 4C). Moreover, we found that priming activity can be rescued at the downstream serine in a ‘priming deficient’ S662A mutant by simply substituting a valine for the +1 leucine at the 2^nd^ phosphorylation site (i.e., FASP S662A/L666V, Figure 5D). Taken together, these results indicate that priming of FASP is highly sensitive to the presence of a valine at the +1 position. Interestingly, the hydrophobic pocket occupied by this valine is created between Tyr^225^ and Leu^173^ of CK1 only when the activation loop is down. The +1 hydrophobic pocket is also small and likely contributes to substrate selectivity based on steric occlusion of bulkier residues at the +1 position, considering that co-crystal structures of CK1 bound to phosphorylated Tap63α PAD and FASP peptides are comprised entirely of backbone-backbone interactions between peptide and kinase in this region (Figure 5E).

We further tested our FASP binding model by introducing an alanine mutation to CK1 at Asp^132^ (D132A) because this residue frequently interacted with the −3 lysine of FASP over the course of the molecular dynamics trajectories. Titration of the ‘priming only’ FASP peptide (S665A) comparing CK1 WT and D132A showed a dramatic loss of kinase activity (Figure 5F). Since Asp^132^ is located directly under the nucleotide binding site and potentially in position to contact ATP, we also sought to test whether the D132A mutation would disrupt kinase activity on a primed FASP peptide, where CK1 activity is driven by the consensus recognition mechanism involving the 1^st^ anion binding site. While the D132A mutation showed a dramatic loss in activity on the unprimed FASP peptide, it had a more modest effect on the primed FASP peptide (Figure S15B-C), suggesting that this mutation primarily reduces activity on FASP via recognition of the unprimed substrate.

One PER homolog within the circadian system, PER3, does not appear to be a substrate of the kinase and lacks the critical residues that precede the priming serine [100]. Here, we further tested our model by introducing a lysine at the −3 position and a glutamate at the −1 position of PER3 FASP-like peptides (A610K/L612E) to mimic the PER2 FASP priming region and rescue priming activity. These mutant peptides were used in a ^32^P-ATP timecourse kinase assay and showed an increase in CK1 activity for both the WT (Figure S15E) and the ‘priming only’ (Figure S15F) substrates in the presence of a −3K and −1E, further demonstrating that these residues play an important role in CK1 recognition and activity on FASP substrates.

### Mapping of binding pockets on the CK1 surface

Our MSMs support that inversion of substrate selectivity in the *tau* mutant is achieved by inversion o the conformational equilibrium in the activation loop, with destabilization of the ‘loop down’ conformation in favor of the alternative ‘loop up’. To understand how the activation loop re-shapes the molecular surface of CK1 and to look for potential allosteric sites controlling substrate selectivity, we screened the CK1 surface using FTMap [101]. For more details, see Supplementary Information (Section E).

#### The activation loop significantly re-shapes the substrate binding cleft in CK1

For both the WT and *tau* mutant, the top-ranked pockets identified by FTMap correspond to the ATP binding site and the Mg^2+^ pocket near the catalytic site (Tables S7 – S10). Interestingly, we observed a fragmentation of the active site and substrate binding cleft in the *tau* mutant, which break down into three separate sub-pockets not as well ranked as the large contiguous pocket detected in the WT enzyme (Figures S16 and S17). Comparison of these pockets when CK1 is in ‘up’ or ‘down’ conformation reveal how dramatically the activation loop conformation re-shapes the substrate binding cleft (Figure 6). The ‘loop down’ conformation promotes a straight binding cleft running contiguously from the first to the second anion binding site, above the activation loop (Figure 6A). The ‘loop up’ conformation, however, appears to promote a bent binding cleft, with part of the substrate channel in the space between helices αD and αF, below the activation loop (Figure 6B). Considering that the WT CK1 has a higher preference for FASP, we hypothesized that the straight binding cleft produced by the more common ‘loop down’ conformation is well suited to bind the FASP substrate, in agreement with our recent co-crystal structures of WT CK1 bound to phosphorylated FASP peptides (pFASP) (Figure 6C) [41]. The bent binding cleft produced by ‘loop up’ conformations could more favorably interact with the pD substrate, which could explain how *tau* not only reduces activity on FASP but also increases the activity on pD substrates. Interestingly, we found that a *Drosophila* PER (dPER) peptide phosphorylated at S589 (perShort peptide), the site of the *per*^S^ mutation that destabilizes dPER and shortens circadian period [102], binds to CK1 when the substrate binding cleft is bent [41]. pS589 of the perShort peptide coordinates the first anion binding site identically to pFASP but follows a channel exposed by the ‘loop up’ conformation of the activation loop (Figure 6D), in excellent agreement with the FTMap analysis.

**Figure 6.**
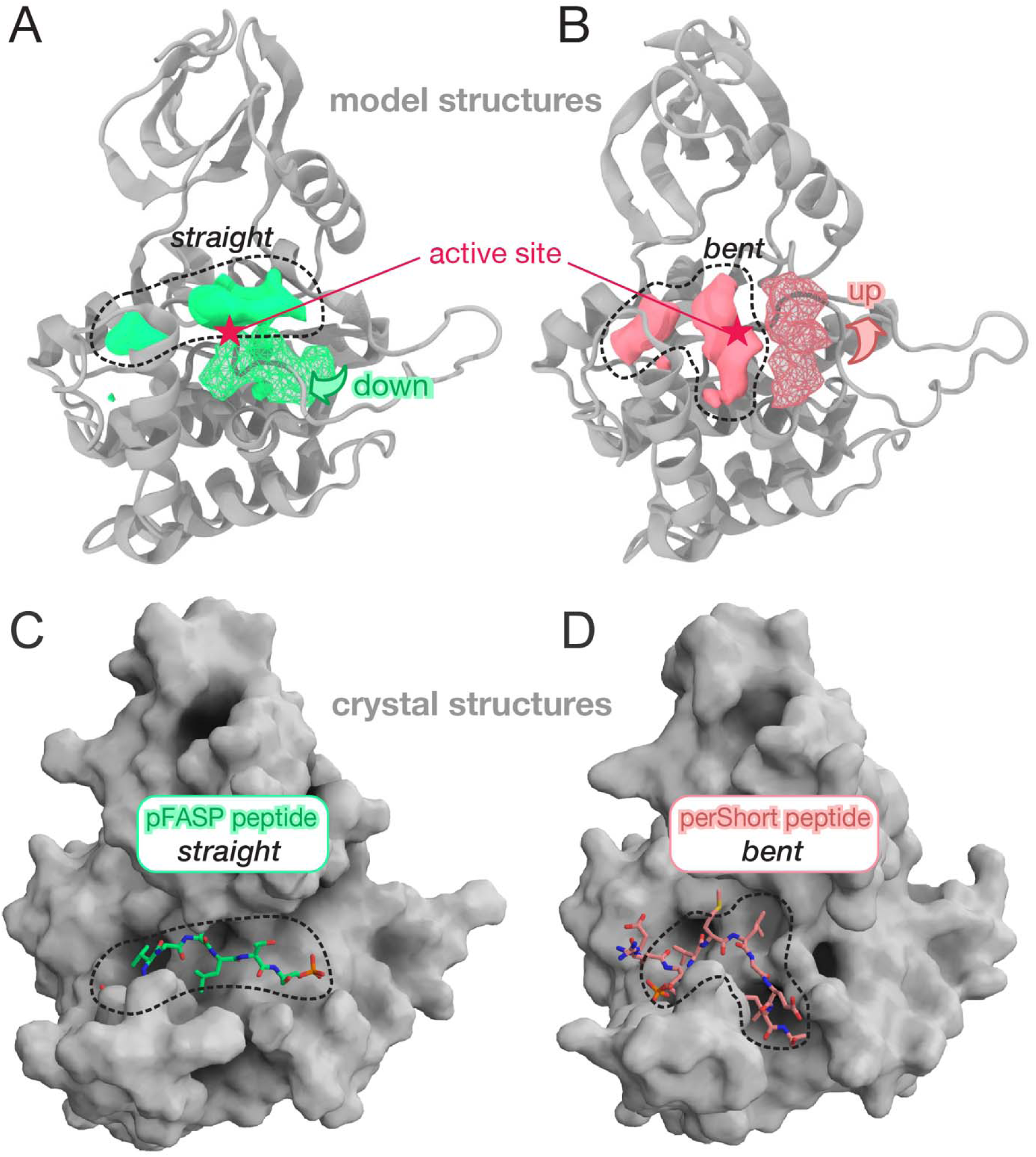
The conformation of the activation loop shapes the substrate binding cleft. A) The ‘loop down’ conformation of the activation loop creates a straight binding cleft whereas (B) the ‘loop up’ conformation creates a bent binding cleft, opening a sub-pocket right under the active site. The solid blobs represent density maps computed based on organic probes from FTMap while the meshed blobs represent occupancy maps computed for residues Lys^171^, Asn^172^ and Leu^173^ along the simulations. C-D) Co-crystal structure of human PER2 pFASP bound to CK1 (PDB 8D7M) in the straight substrate binding cleft (C), whereas the dPER perShort peptide binds to CK1 (PDB 8D7P) following a bent substrate binding cleft (D).

#### An allosteric pocket to shorten the circadian rhythm

FTMap also identified potential allosteric sites on the surface of CK1, as described in Supplementary Information (Section E). Of particular interest is a pocket formed between helix αC and the activation loop (Figure 7A). As highlighted in Figure 7A, this “activation” pocket includes three residues belonging to the activation loop (Lys^171^, Asn^172^ and Leu^173^) and it is only fully assembled when the activation loop is up. Indeed, this pocket is ranked higher in the *tau* mutant (Table S10) than in the WT enzyme (Table S9), in agreement with the fact that *tau* stabilizes the upward conformation of the activation loop. Other key residues forming this potential allosteric pocket are His^46^ and Pro^47^ located in the loop connecting sheet β3 to helix αC. Interestingly, mutations at these positions (H46R and P47S) of the *Drosophila* CK1 ortholog, *Doubletime,* shorten the circadian period by ∼ 4 hours (Figure 7B) [50]. Thus, occupation of this allosteric pocket by small molecules mimicking the sidechains of serine and arginine could increase CK1 activity against the pD by stabilizing the ‘loop up’ conformation and bending the substrate binding cleft.

**Figure 7.**
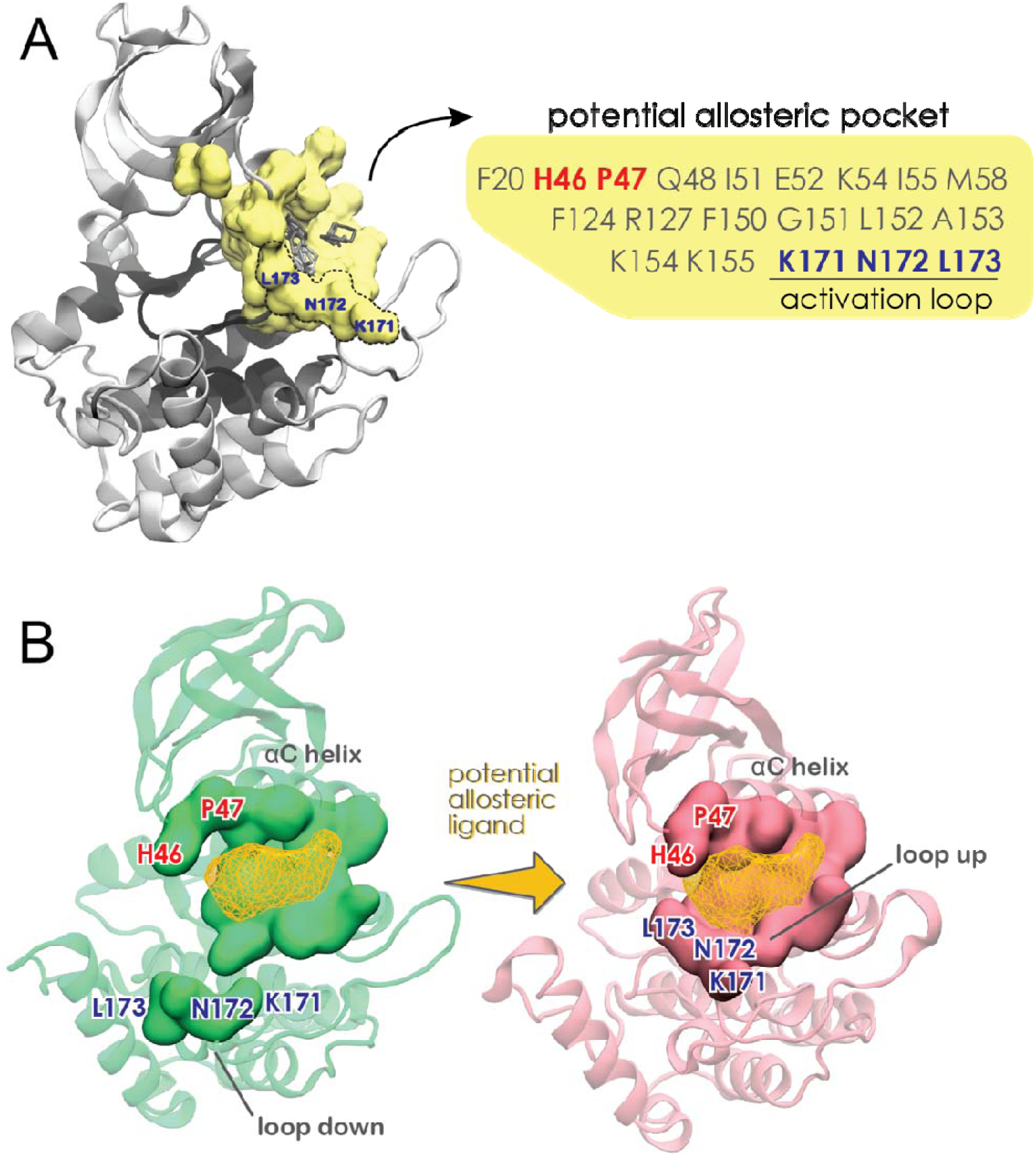
A potential allosteric pocket to shorten the circadian rhythm. A) Residues belonging to the *tau* activation pocket are highlighted in yellow, while the small organic molecules used as probes by FTMap are represented as grey sticks. B) Complete assembly of this pocket occurs when the activation loop is ‘up’. Ligand binding to this pocket is likely to stabilize the ‘loop up’ conformation.

## Discussion

Despite its central role in governing circadian rhythms, the molecular mechanism underlying CK selectivity for its target regions of the circadian protein PER remains poorly understood, hindering rational attempts to develop CK1-specific circadian drugs. Because CK1 is generally thought to be constitutively active towards primed or acidic substrates [62, 66–68, 70], the role of its activation loop has been largely unexplored. In this study, we use an extensive set of molecular dynamics simulations integrated into Markov State Models to map the free energy landscape governing the slow conformational transitions of CK1’s activation loop. Our model reveals the existence of a slow equilibrium between the more stable ‘loop down’ and the less stable ‘loop up’ conformations in the wild-type enzyme and shows that the period-shortening *tau* mutant biases and accelerates the transitions towards the ‘up’ conformation.

We also found that the dynamics of the activation loop are strongly connected to the conformational state of loop EF and that flipping of the activation loop requires transient unfolding of loop EF. This explains the long timescales associated with ‘up-down’ transitions, which would happen faster if the conformation of the activation loop was controlled mainly by the configuration of Gly^175^, as previously hypothesized [71]. Elucidation of the role of Gly^175^ for the conformational dynamics of the activation loop reinforces the importance of molecular dynamics in re-visiting and adding complexity to structural hypotheses that are based on static observations. These findings also explain how the R178C *tau* mutation, which facilitates the unfolding of loop EF by disrupting the first anion binding site, significantly accelerates the dynamics of the activation loop. Interestingly, given the importance of loop EF for temperature-compensation against circadian substrates [31, 66, 70] and its allosteric connection with the activation loop [41], it is likely that the dynamic equilibrium involving the activation loop is also part of CK1 temperature-compensation mechanisms.

This work characterizes for the first time how CK1 interacts with the unprimed PER FASP region. Our model of the FASP interaction with CK1 suggests key interactions important for the non-consensus priming step that are supported by *in vitro* biochemistry. In agreement with the relatively low atomic fluctuations of the +1 residue (relative to the priming site) from our model, our data clearly demonstrate the importance of the +1 valine (+1V) for non-consensus priming of FASP, which fits into a small hydrophobic pocket created between Leu^173^ and Tyr^225^ on CK1. Our model also reveals that FASP makes use of the first anion binding site by partially inserting a negatively charged glutamate at position −1 (−1E) of the substrate. Additionally, we showed that stabilization of FASP in the binding cleft is complemented by electrostatic interactions between the lysine at position −3 (−3K) with Asp^132^ of CK1 as well as the β-phosphate group of ATP. Finally, we demonstrated that addition of a −3K and −1E to PER3 was sufficient to allow phosphorylation of the priming serine by CK1. Together these findings establish −3K, −1E, and +1V as bona fide residues necessary for proper CK1 recognition of FASP as a substrate.

Interestingly, while the −3K-ATP interaction can lock the FASP substrate in place for the priming event, we speculate that this interaction could also facilitate unbinding (or translocation) of the primed product by attaching it to the leaving ADP sub-product. In addition, substrate stabilization achieved by partial occupation of the first anion site by −1E could be just enough to promote priming without excessive stabilization of the primed product. In consensus substrates, this anion site is fully occupied by a negatively/phosphorylated residue at position −3, as recently shown in the structures of CK1 bound to primed Tap63a or PER FASP peptides [41, 72]. It can even be double occupied (by phosphorylated serines at positions −3 and −6) in the case of multiply phosphorylated Tap63α 72], or as seen in one crystal structures of CK1 in anionic solvent conditions [31]. This suggests that the partial occupation of the first anion site by unprimed FASP is likely to be replaced with progressively stronger occupation of this site by phosphorylated serine at positions −3 to allow subsequent phosphorylation events. In a processive or semi-processive mechanism, these escalating electrostatic interactions at the upstream region of FASP would provide a direction for the sequential phosphorylation events, while the lack of specific interactions in the downstream region would make it easier for the next phosphorylation site to translocate into the active site.

Using FTMap to map the surface of CK1 in different conformations, we demonstrated how the conformation of the activation loop re-shapes the substrate binding cleft. Applying FTMap to the conformations extracted from our MSM also allowed us to identify a potentially attractive allosteric site between helix aC and the activation loop. Interestingly, this site partially overlaps with the site that accommodates a phosphate group in kinases that are activated by phosphorylation of the activation loop [71], suggesting a possible regulatory role. To add to that, two period-shortening mutations are found in this site: P47S and H46R. Differently from *tau* both P47S and H46R mutants achieve their period-shortening effects by increasing activity on the pD region while retaining normal FASP priming activity relative to WT [71]. This is consistent with what would be expected from increasing the population of ‘loop up’ and ‘bent’ substrate binding cleft conformations (which favor pD phosphorylation) without disruption of the 1^st^ anion binding pocket (important for priming and subsequent phosphorylation of FASP). Because this pocket is only fully assembled when the activation loop is ‘up’, we propose that it might be a viable site to pharmacologically stabilize the ‘up’ conformation and shorten the period of circadian rhythms.

## Author contributions

C.G.R., J.M.P. designed research, performed experiments, analyzed data, and wrote the manuscript. M.R.T. prepared materials, performed experiments and edited the manuscript. A.M.F., R.N. prepared materials, performed experiments, and analyzed data. E.P. performed experiments and analyzed data. R.A., D.M.V., A.M., C.L.P. guided research, funded research, and edited the manuscript.

## Acknowledgements

Funding for this work was provided by the US National Institutes of Health grant R35 GM141849 (C.L.P.). D.M.V. was supported by Singapore Ministry of Health grant MOH-000600. J.A.M. was supported by TSCC computer resources from UC San Diego. C.L.P. was supported by the Howard Hughes Medical Institute.

## Supporting materials

Supporting materials and additional methods can be found online.

