## Supplemental Materials and Methods for "Markovian State Models uncover Casein Kinase 1 dynamics that govern circadian period"

#### A. Molecular dynamics simulations of apo proteins

To start the molecular dynamics simulations of WT and *tau* CK1 that would be used to build the MSMs, we extracted conformations from Gaussian accelerated MD simulations as described previously [1] and illustrated in Figure S1. We wanted these initial structures to capture different conformations of the activation loop, so we applied principal component analysis (PCA) to the backbone atoms of the activation segment and its flanking regions (residues 151 to 185, Figure S1A). All the GaMD trajectories from WT or *tau* CK1 were concatenated together so that the resulting vectorial space (PC1 vs PC2) is the same for both systems. We then selected 10 conformations located at different regions of the essential space describing the activation loop dynamics and used them as starting points for our first round of conventional MD simulations (Figure S1B). For each initial structure, we launched three independent MD replicas (with different initial velocities) for 300 ns, totalizing 9us for each system (WT or *tau*). After these finished running, we randomly selected 20 more structures from the new trajectories and re-launched two independent MD replicas for 300 ns, adding 12us to the total time simulated for each system.

All systems were solvated with TIP3P water molecules [2] in cubic boxes with at least 15 Angstroms between the protein and the box boundaries. The simulation boxes were neutralized with Na+ or Cl- counterions. Protein and ions were parametrized with Amber ff14SB forcefield [3], while parameters for $SO_4^{2+}$ (the anion bound to the first anion binding pocket in the WT CK1) were obtained from the Generalized Amber Force Field (GAFF) [4] and adjusted as previously proposed [5].

Conventional MD simulations were performed with AMBER 16 [6] in the NVT regime, with a time step of 2fs. The PME method [7] was used to calculate electrostatic interactions using periodic boundary conditions. A 12 Angstroms cutoff was used to truncate non-bonded short-range interactions.

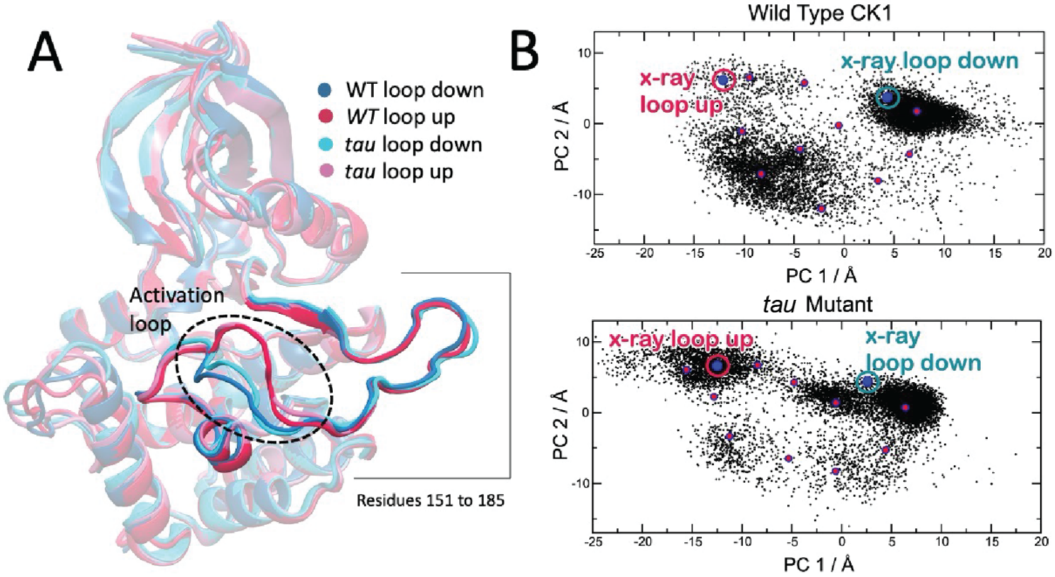


Figure S1. Selection of conformations from previous GaMD simulations as starting points for the MD simulations used to generate MSMs. A) Region of CK1 selected for Principal Component Analysis. B) Conformations from GaMD simulations selected for starting conventional simulations (pink dots).

### B. Markovian State Models (MSMs)

#### Main (distance-based) model

Before building the MSMs, we used root mean square fluctuation (RMSF) profiles to identify the most flexible or rigid regions of the proteins (Figure S2A). We then selected 9 pair-wise distances involving the activation loop (Leu173) and the C-terminal portion of L-EF (T220). All these distances involved functionally important residues of protein, some of them flexible (S19 at the Gly-rich loop and T44 at the L-3A loop) and some of them rigid (D128 in the HRD motif and G151 in the DFG motif) to provide good anchor points. Pair-wise distances between these residues were calculated based on the positions of the α-carbons accumulated over WT and *tau* trajectories and used as input features for time-lagged Independent Component Analysis (tICA) [8]. tICA essentially identifies the linear combination of features that best describes the *slowest* modes of motions of the system. The resulting orthogonal components (or TICs) are ranked by their associated implied timescales (from slowest to fastest). Figure S2B shows that the slowest mode of motion (TIC 1) consistently involves Leu173 at the activation loop, while the second slowest motion (TIC 2) consistently involves T220 at L-EF. In particular, the feature that most contributes to TIC 1 is the distance between Leu173, in the activation loop, and G151, which is part of the DFG motif, a rigid region of the protein. These results show that rearrangement of the activation loop is the slowest mode of motion in our model, even though L-EF displays higher amplitude of motion as measured by atomic fluctuations (RMSF). We finally selected the five distances listed in Figure S2C to construct our main MSM.

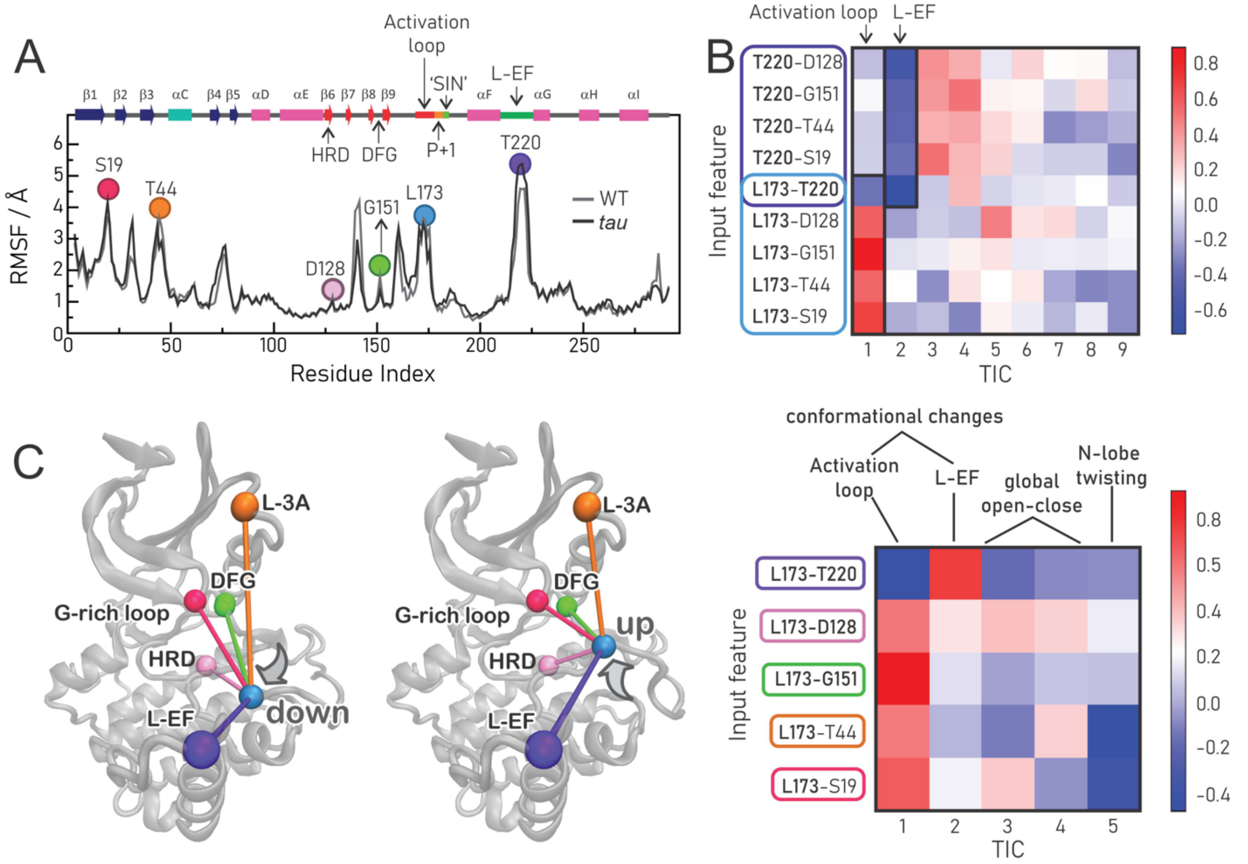


Figure S2. Selection of distances for the MSM model. A) Backbone RMSF with respect to average structure for concatenated WT or *tau* trajectories. Alignment was performed with the rigid core of the enzyme (residues 85 to 165). The residues whose pair wise distances were selected as input features are highlighted. B) tICA results showing the correlation of input features with each TIC component (1$^{st}$ and 2$^{nd}$ are highlighted). C) Final pair wise distances selected for tICA and MSM construction.

We used PyEmma [9] version 2.5.7 to process the trajectories and to build, validate and analyze the MSMs. Discretization of the conformational space was performed for WT and *tau* separately using k-means clustering, with k = 200 for both systems. The corresponding implied timescale plots are shown in Figure S3A. We then used Hidden Markov Models to coarse-grain the MSMs into fewer metastable states that are more amenable to human interpretation. Metastable states represent kinetically distinct conformations separated by high-energy barriers (or slow motions). We decided on the number of metastable states for each system based on spectral analysis of the tICA modes of motion (Figure S3B) [10, 11]. For both WT and *tau*, we observed a spectral gap between the 2$^{nd}$ and 3$^{rd}$ timescales, indicating the dominance of two slow modes of motion and three well separated minima in the conformational landscape, so we coarse-grained the MSM with three metastable states for both systems. Clustering of microstates into metastable states was performed with the PCCA++ algorithm [12-14]. The results are shown in Figure S4A, along with the visualization of the 2$^{nd}$ and 3$^{rd}$ eigenvectors of the transition matrix, which represent the two slowest modes of motion (the 1$^{st}$ eigenvector corresponds to the equilibrium distribution) (Figure S4B). We then constructed Bayesian Hidden Markov Models (HMSM) [15, 16], with n = 3 and lag time ($\tau$) of 3 ns. The quality of the coarse-grained HMSM was verified by ITS plots (Figure S5A) and Chapman-Kolmogorov (CK) tests (Figure S5B,C), with 95% confidence intervals calculated using a Bayesian sampling scheme. These are the models presented and discussed in the main manuscript.

To characterize the conformations trapped in each metastable state, we extracted 5000 representative structures with at least 70% of assigned membership. To describe the activation loop, we measured the α–carbon distance between Leu173, in the activation loop, and Leu152 (located at a rigid region of the protein) (Figure S6A). To characterize the conformational state of L-EF, we measured the RMSD of this region with respect to the crystallographic structure (Figure S67). Based on this analysis we classified the metastable states as described in Table S1. Mean first passage times (MFPTs) between states are reported in Table S2.

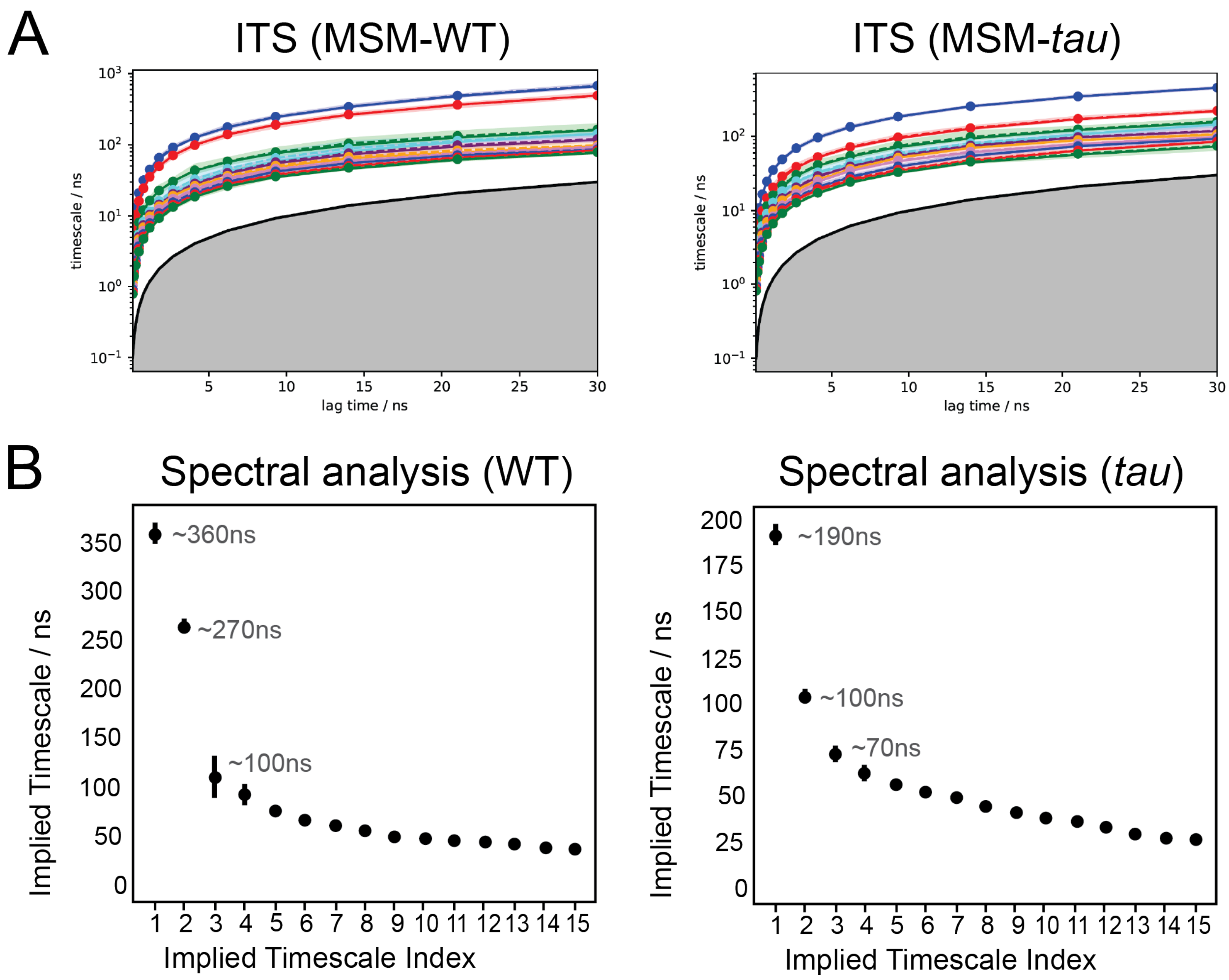


Figure S3. Analysis of the MSM model for coarse-graining. A) Convergence of MSM implied timescale plots. The grey area marks the lower limit of timescales that can be resolved for each lag time. Colored shaded areas represent 95% confidence intervals estimates from Bayesian-sampled transition matrices. Dashed lines represent the mean of Bayesian MSMs. B) Spectral analysis of implied timescales.

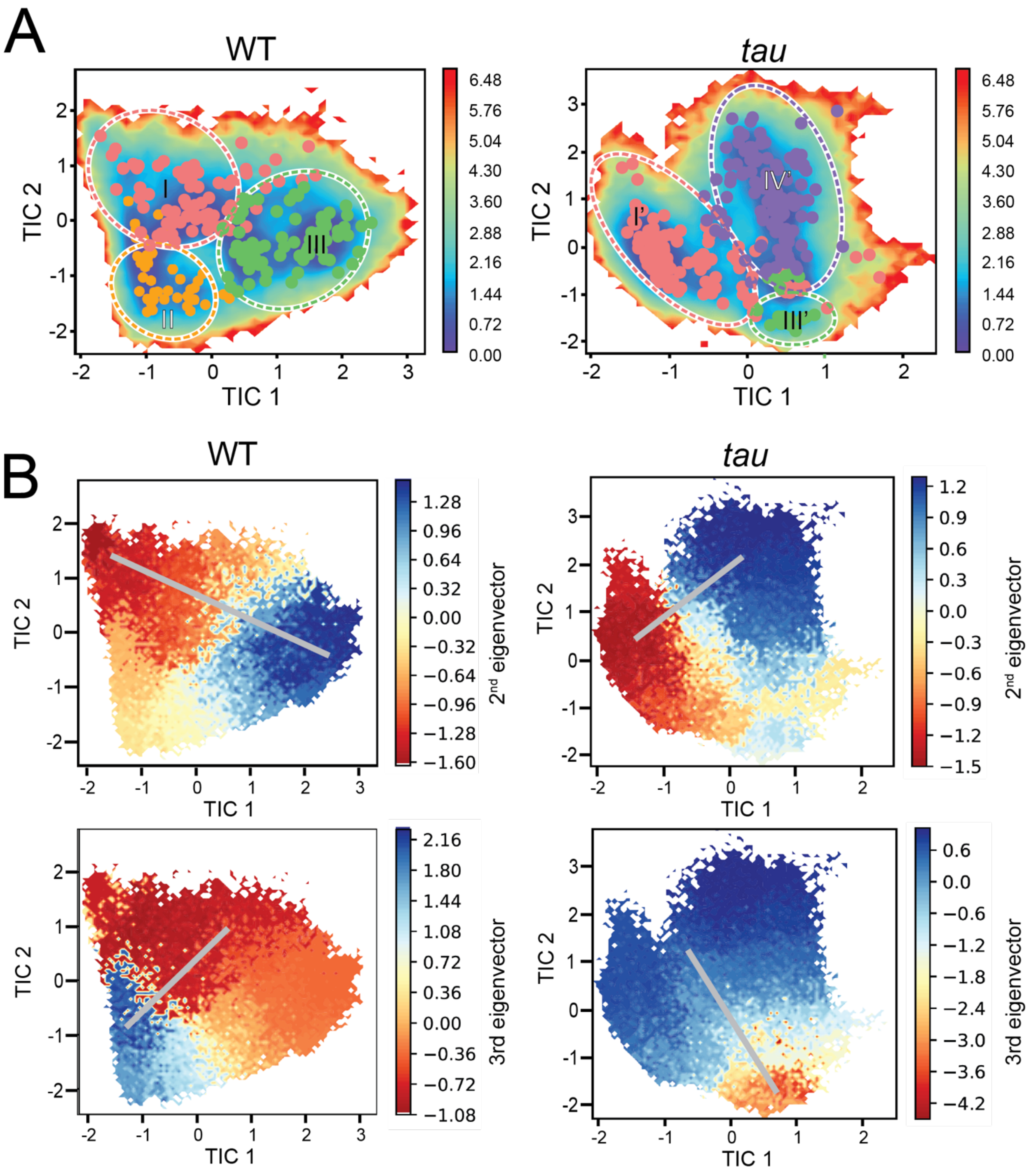


Figure S4. Analysis of metastable states. A) Free energy landscapes (FEL) superimposed with microstate clusters assigned to metastable states (I/I', II, III/III', and IV'). FELs were obtained by re-reweighting the trajectory frames with the stationary probability distribution (1st eigenvector) derived from the MSM transition matrix. B) Probability shifts associated with the 2nd and 3rd eigenvectors, which represent the 1st and 2nd slowest conformational changes in the system.

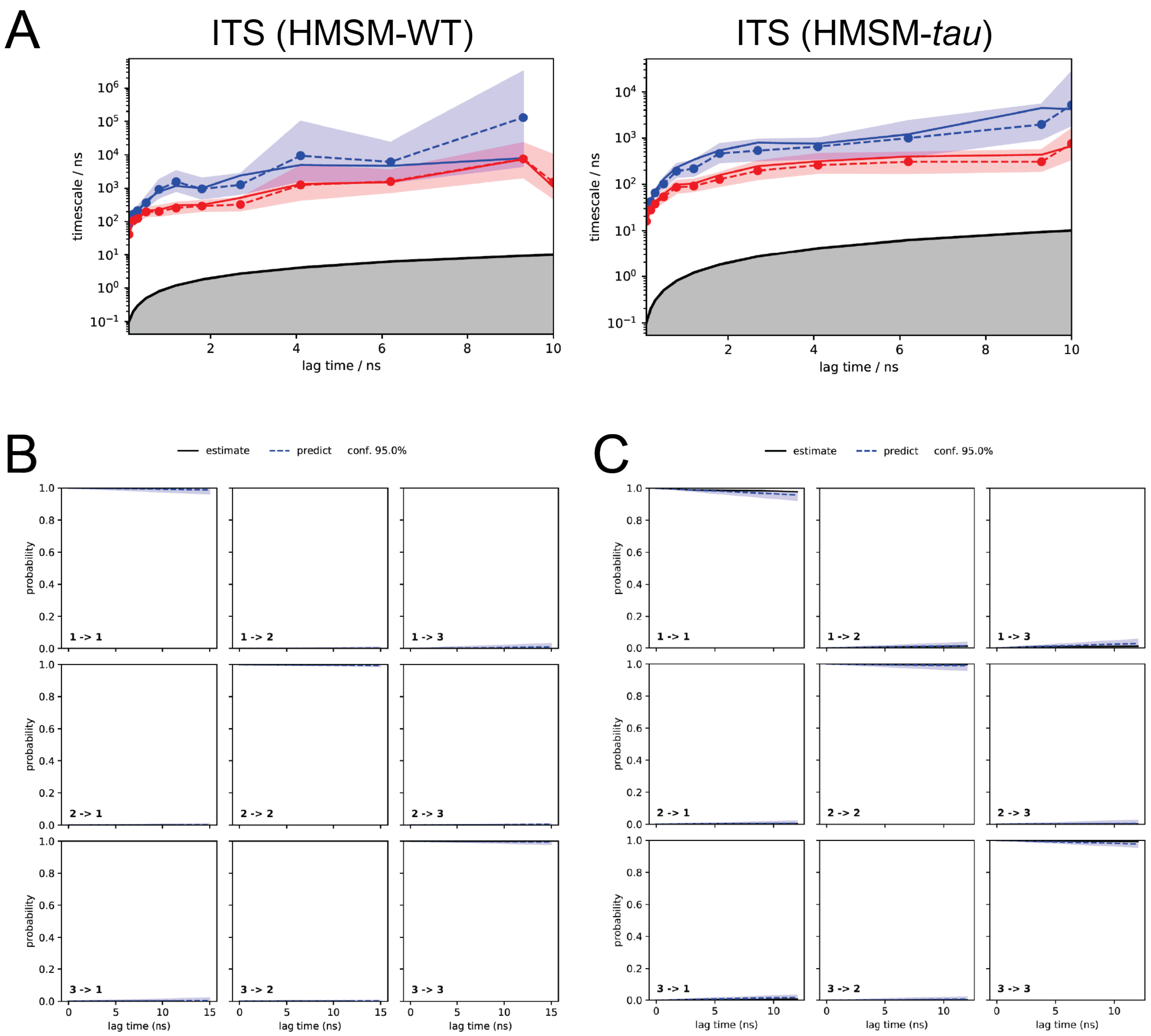


Figure S5. Quality assessment of the model. A) Convergence of ITS plots for coarse-grain HMSM. B,C) Chapman-Kolmogorov (CK) tests for HMSMs obtained for WT CK1 (B) and *tau* mutant (C). Shaded areas in the plots represent 95% confidence intervals estimated from Bayesian-sampled transition matrices and dashed lines represent the mean of Bayesian MSMs.

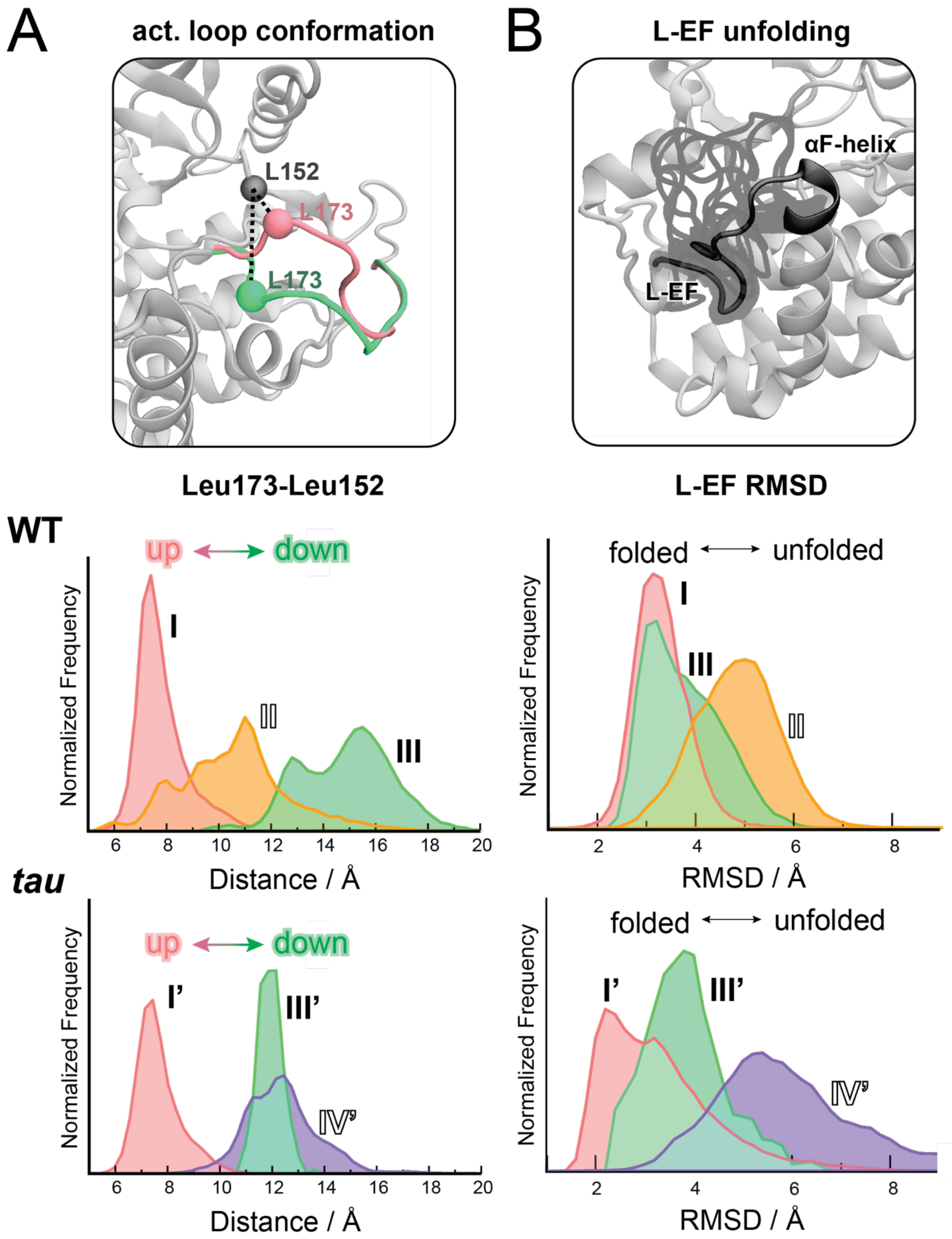


Figure S6. Characterization of meta-stable states in terms of the conformation of the activation loop. A) The conformation of the activation loop (cα distance between Leu173 and Leu152) and (B) conformational state of L-EF (high RMSD values indicate unfolding) for WT (top) and *tau* mutant (bottom). Histograms are normalized and do not reflect the relative populations of each state. Distances were calculated between the γ-carbon atom of Leu[173] and the α-carbon of Leu[152]. RMSDs were calculated for the backbone of residues 212 to 226 (loop EF and top of helix F), after alignment was performed for residues 85 to 165 with respect to the average structure.

Table S1. Classification of main metastable states

| Metastable state | % population in WT | % population in *tau* | Conformation of the activation loop | L-EF conformational state |
|---|---|---|---|---|
| I/I’ | 34 +- 25 | 48 +- 9 | Up | Folded |
| II | 19 +- 14 | --- | Intermediate | Unfolded |
| III/III’ | 47 +- 21 | 20 +- 6 | Down | Folded |
| IV’ | --- | 32 +- 8 | Down | Unfolded |

Table S2. MFPT between metastable states from main HMSM

| Transitions in WT | MFPT / µs | Transitions in *tau* | MFPT / µs |
|---|---|---|---|
| I → II | 10.9 | I’ → III’ | 1.6 |
| II → I | 13.9 | III’ → I’ | 1.2 |
| I → III | 8.8 | I’ → IV’ | 1.7 |
| III → I | 14.9 | IV’ → I’ | 1.4 |
| II → III | 3.3 | III’ → IV’ | 0.9 |
| III → II | 6.5 | IV’ → III’ | 0.9 |

#### Gly$^{175}$ (torsional) model

The second MSM model was constructed with input features based solely on the backbone angles of Gly$^{175}$ (Figure S7A) since this residue was hypothesized to work as a hinge controlling the conformation of the activation loop [1]. Discretization of Gly$^{175}$ configurational space was performed for WT and *tau* separately, using k-means clustering with k = 200 for both systems. Converged ITS plots revealed a gap between the 1$^{st}$ and 2$^{nd}$ timescales in WT and between the 2$^{nd}$ and 3$^{rd}$ timescales in the *tau* mutant (Figure S7B). We thus clustered the microstates into 2 metastable states for the WT and 3 metastable states for the *tau* mutant (Figure S7C). HMSM were generated with τ=20ns and validated as shown in Figure S7 (panels D-F).

The resulting HMSMs are displayed in Figure S8A and detailed in Tables S3 and S4. Conformational characterization of metastable states (Figure S8B) reveal that the torsional models fail to separate the two conformations of the activation loop. This is especially the case of WT, where metastable states A and B contain both ‘down’ and ‘up’ conformations within the same state. The difference between these two states is that in state A the activation loop can also adopt intermediate conformations between ‘up’ and ‘down’. For the *tau* mutant, the torsional model performs slightly better: while the most populated state (B’) still mixes ‘up’ and ‘down’ conformations, states C’ and D’ consist of (mainly) ‘loop up’ and ‘loop down’ conformations, respectively.

We next plotted representative structures of the metastable states in the Ramachandran space of Gly$^{175}$ (Figure S9A). To investigate whether the configuration of Gly$^{175}$ is correlated with the conformation of the activation loop, we also colored the Ramachandran plots according to the distance between Leu$^{173}$ and Leu$^{152}$ (Figure S9B). Visual inspection of these plots reveals no clear correlations in the WT but suggests a moderate correlation between the conformation of the activation loop and $\phi^{Gly175}$ in *tau*. To verify this, we performed a linear regression between $\phi^{Gly175}$ and the distance between $Leu^{173}$ and $Leu^{152}$, as shown in Figure S9C. The resulting correlation coefficients confirm a weak correlation between the $\phi^{Gly175}$ and the conformation of the activation loop in the *tau* mutant (r=0.5), with negative values of $\phi^{Gly175}$ favoring 'loop up' conformations and positive values of $\phi^{Gly175}$ favoring 'loop down' conformations. In the WT enzyme, this correlation is not significant (r=0.36).

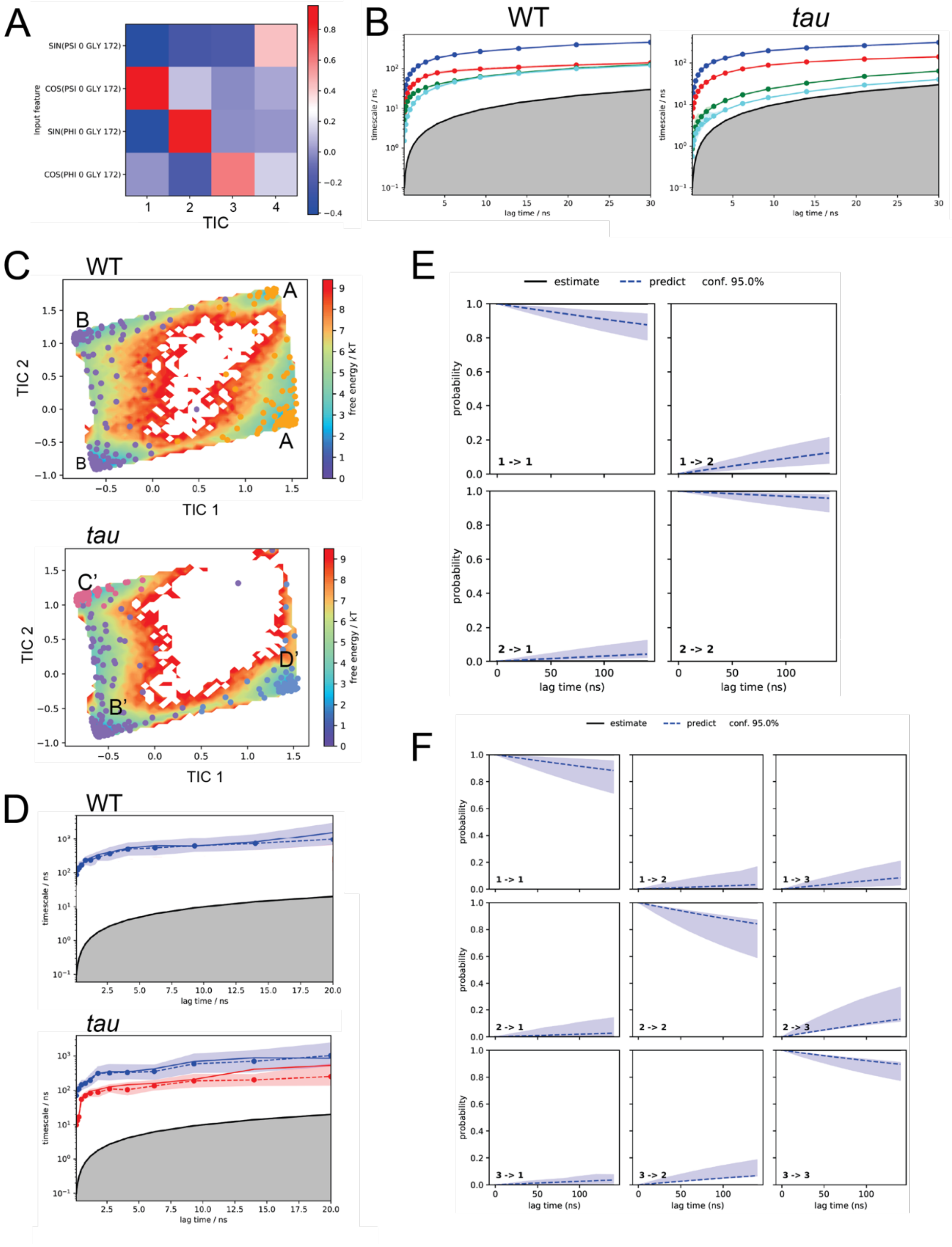


Figure S7. MSM model based on Gly$^{175}$ torsions. A) tICA results using Gly$^{175}$ backbone angles as input features. B) Convergence of MSM implied timescale plots. The grey area marks the lower limit of timescales that can be resolved for each lag time. Colored shaded areas represent 95% confidence intervals estimates from Bayesian-sampled transition matrices. Dashed lines represent the mean of Bayesian MSMs. C) Resulting free energy landscapes in terms of the slowest tICA components, with microstates clustered into meta-stable states (identified by letters). D) Convergence of ITS plots for coarse-grain HMSM. E,F) Chapman-Kolmogorov (CK) tests for HMSMs obtained for WT CK1 (E) and *tau* mutant (F). Shaded areas in the plots represent 95% confidence intervals estimated from Bayesian-sampled transition matrices and dashed lines represent the mean of Bayesian MSMs.

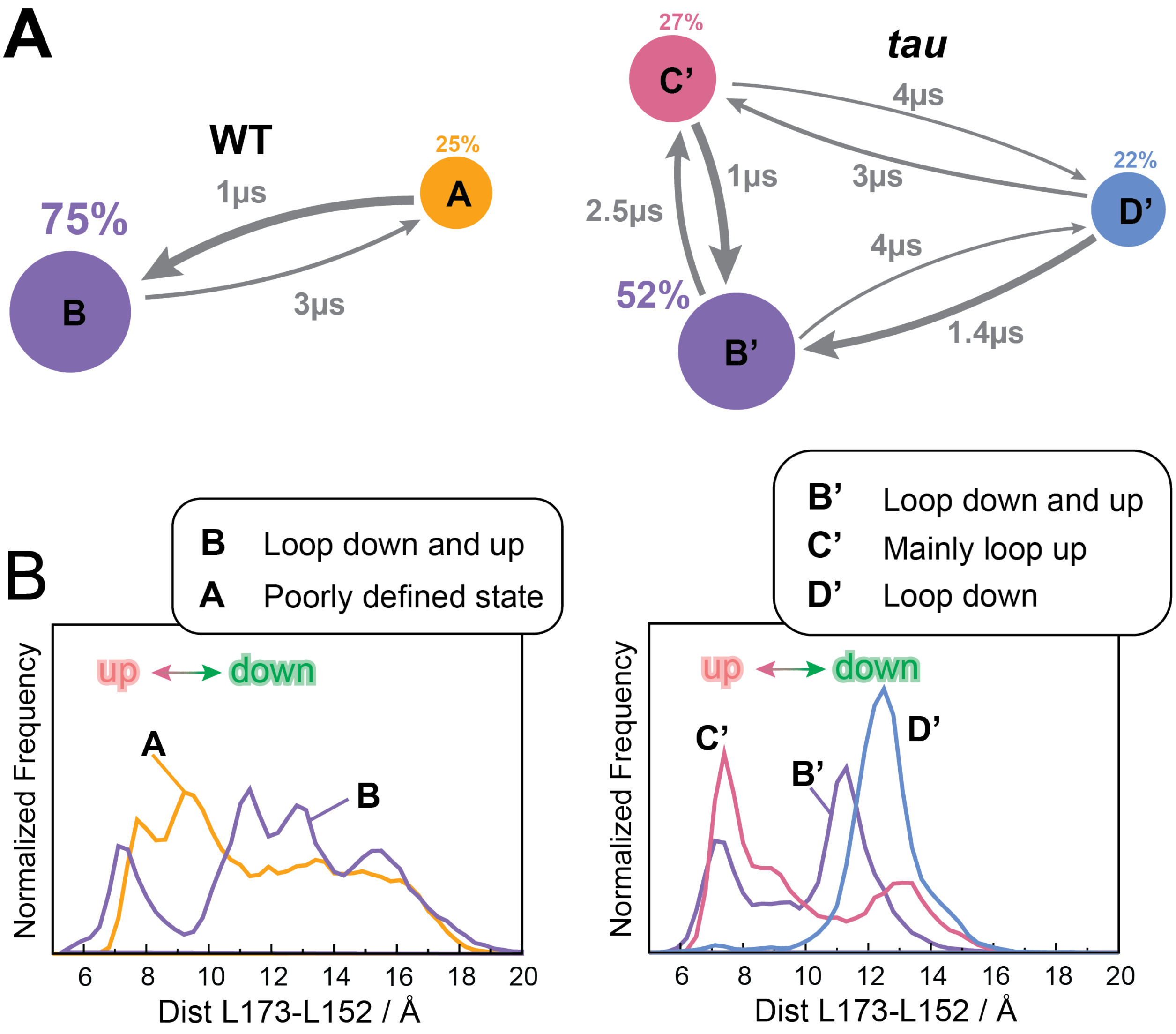


Figure S8. Characterization of the $Gly^{175}$ MSM model in terms of the conformation of the activation loop. A) Equilibrium populations of meta-stable states and MFPTs between states for the WT (left) and *tau* (right) protein systems. Area of the circles is proportional to the equilibrium population and thickness of the arrows is proportional to transition rates between states. The numbers next to the arrows indicate MFPTs. B) Characterization of meta-stable states in terms of the conformation of the activation loop (α-carbon distance between $Leu^{173}$ and $Leu^{152}$) for the WT (left) and *tau* (right) protein systems. Histograms are normalized and do not reflect the relative populations of each state. Distances were calculating between the γ-carbon atom of $Leu^{173}$ and the α-carbon of $Leu^{152}$.

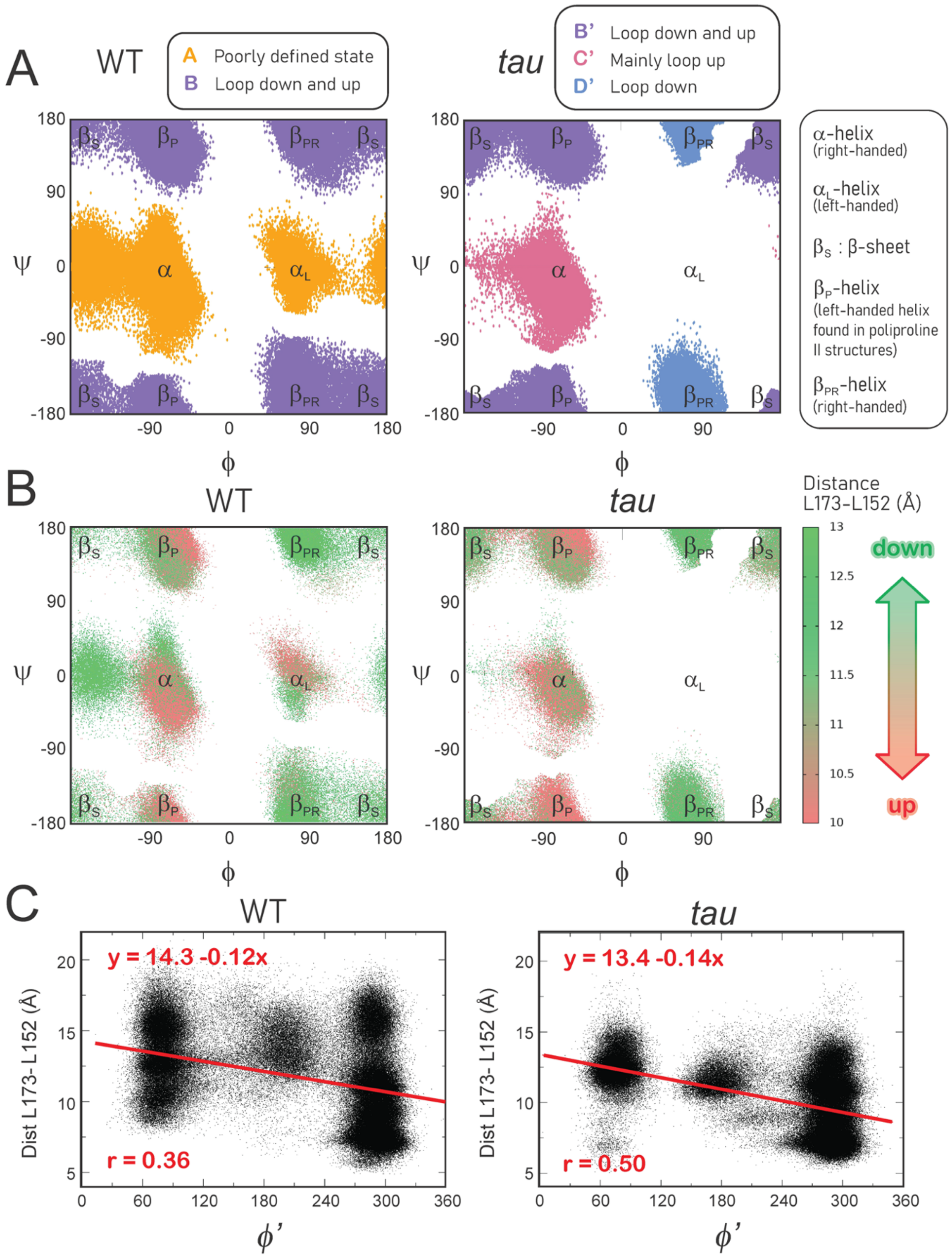


Figure S9. Meta-stable states produced by the Gly[175] MSM model do not correlated with the conformation of the activation loop. A) Gly[175] Ramachandran space colored according to metastable state assignment for WT (left) and *tau* (right). Low energy regions are indicated as described previously [17]. B) Gly[175] Ramachandran space colored according the (cα) distance between Leu[173] and Asp[152]. C) Linear regression between $\phi^{G175}$ and the (cα) distance between Leu[173] and Leu[152].

Table S3. Classification of torsional HMSM metastable states

| Metastable state | % population in WT | % population in *tau* | Conformation of the activation loop |
|---|---|---|---|
| A | 25 +- 10 | --- | Poorly defined |
| B/B’ | 75 +- 10 | 52 +- 13 | Up and down |
| C | --- | 27 +- 10 | Mainly up |
| D | --- | 21 +- 15 | Down |

Table S4. MFPT between metastable states from torsional HMSM

| Transitions in WT | MFPT / µs | Transitions in *tau* | MFPT / µs |
|---|---|---|---|
| A → B | 1.1 | B’→ C’ | 2.5 |
| B → A | 3.1 | C’→ B’ | 1.2 |
| | | B’→ D’ | 4.1 |
| | | D’→ B’ | 1.4 |
| | | C’→ D’ | 4.3 |
| | | D → C’ | 2.9 |

### C. Molecular models of CK1 bound to FASP peptide

#### Initial model based on TAp63α-CK1 complex

To model the interaction between FASP and CK1, we used the x-ray structure of CK1 bound to a double phosphorylated TAp63α peptide as template (PDB 6RU7, chains A and C) [18]. Alignment between FASP and TAp63α peptide sequences is shown in Figure S10A. In this template, CK1 displays the activation loop in the ‘loop down’ conformation, which is the preferred conformation in the wild-type enzyme according to our MSMs. Considering that FASP is the preferred substrate over the pD, it is reasonable to assume that FASP binds to the most common ‘loop down’ conformation. Moreover, both the FASP priming motif and the TAp63a peptide in the template structure display a valine at position +1, which, in the template structure, interacts with Leu173 in the ‘loop down’ conformation.

To model the interactions prior to the priming event, the original ADP molecule in the template structure was computationally replaced by an ATP molecule extracted from an ATP-bound PKA structure (PDB 1ATP), after the active site of the two kinases were superimposed with Lovoalign [19]. We also removed any crystallographic water and ions and manually added $Mg^{2+}$ ions coordinated to the ATP molecule as shown in Figure S10B. The template was processed with Maestro to have hydrogens added, bond orders assigned, and atomic positions optimized. We then mutated the sidechains one at a time (from N- to C-terminal), minimizing the structure after each mutation, until the bound peptide matched the FASP sequence (Figure S10B).

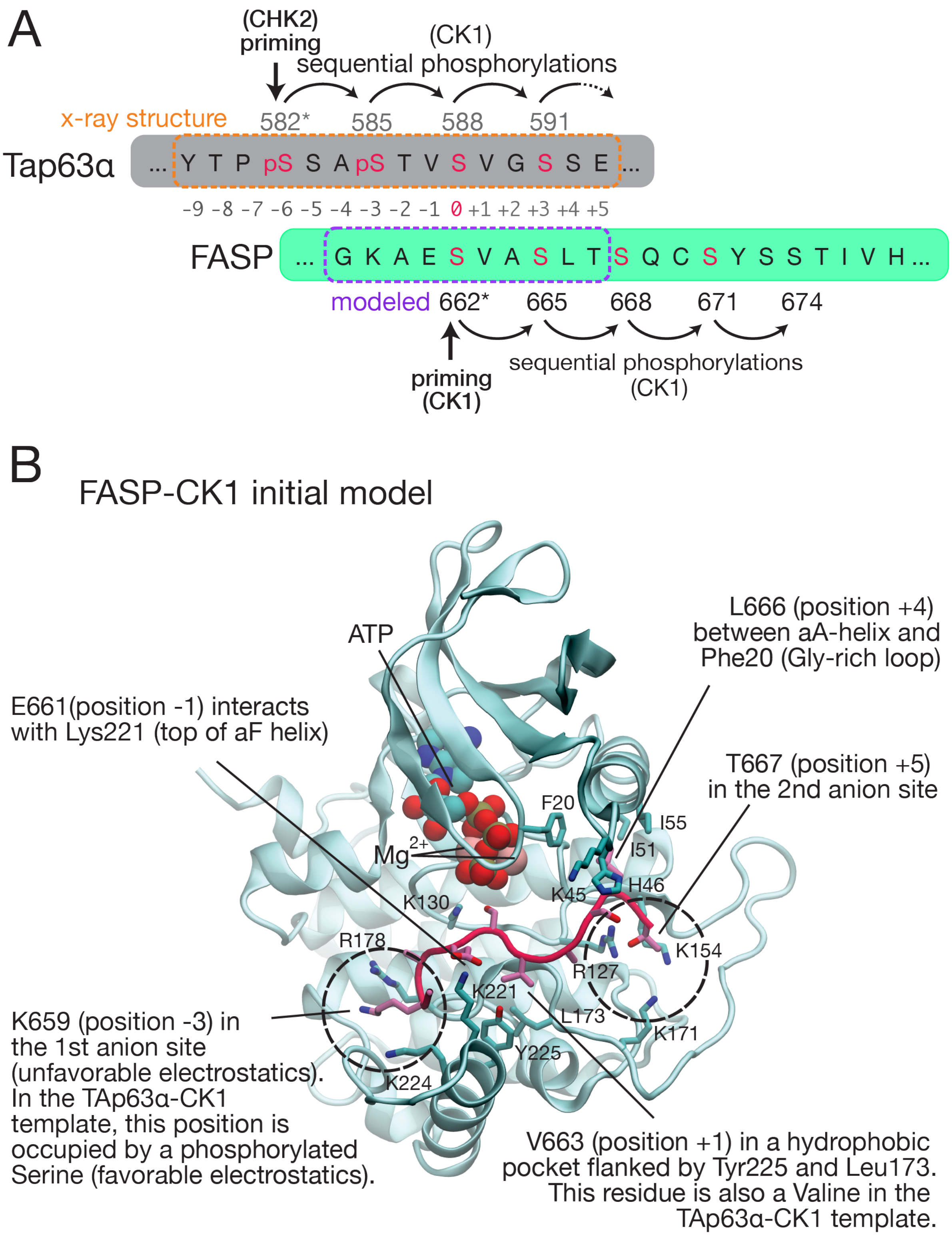


Figure S10. Model used as a starting point for MD simulations A) Alignment between TAp63α peptide and the priming region of human PER2 FASP. B) Molecular model of FASP-CK1 based on PDB 6RU7.

#### Relaxation of the model with Gaussian accelerated MD simulations

To avoid spurious electrostatic interactions during the MD simulations, the N- and C-terminals of the FASP peptide were capped with acetyl (ACE) and amide (NME) capping groups, respectively. Prior to simulations, hydrogen atoms were reassigned with PROPKA [20, 21] at pH 7.0. Parameters for protein residues were obtained from AMBER ff14SB force field [3] while parameters for ATP were obtained as previously described [22].To allow the system to relax and to accelerate the exploration of the binding landscape, we initially ran 10 independent replicas of GaMD simulations [23] starting with different random velocities. Each replica was simulated for 100 ns, totaling 2µs of accelerated simulations. The simulations were performed with AMBER 17 [24] using a time step of 1 fs. Equilibration and simulation protocols are summarized in Table S5.

Table S5. GaMD equilibration and simulation protocol.

| Stage | Position restraints | k (kcal/mol/Å$^2$) | T(K) | P(20) | Steps | Time(ns) |
|---|---|---|---|---|---|---|
| Equilibration | | | | | | |
| Minimization 1 | CK1, FASP, ATP, $Mg^{2+}$ (a.a) | 500 | --- | --- | 2x$10^3$ | --- |
| Minimization 2 | CK1 and FASP (a.a) | 500 | --- | --- | 1x$10^3$ | --- |
| Minimization 3 | CK1 (a.a.) and FASP (bb.) | 500 | --- | --- | 1x$10^3$ | --- |
| Minimization 4 | CK1 and FASP (bb.) | 500 | --- | --- | 1x$10^3$ | --- |
| Minimization 5 | CK1 (bb.) | 500 | --- | --- | 1x$10^3$ | --- |
| Minimization 6 | --- | --- | --- | --- | 1x$10^3$ | --- |
| Thermalization | CK1, FASP, ATP, $Mg^{2+}$ (a.a.) | 30 | 300 | --- | 25x$10^3$ | 0.05 |
| Density equil. | CK1, FASP, ATP, $Mg^{2+}$ (a.a.) | 30 | 300 | 1 | 5x$10^5$ | 1 |
| GaMD preparation | | | | | | |
| cMD prep. | --- | --- | 300 | --- | 2x$10^6$ | 4 |
| GaMD prep. | --- | --- | 300 | --- | 24x$10^6$ | 48 |
| Production | | | | | | |
| GaMD (x10 replicas) | --- | --- | 300 | --- | 1000x$10^6$ | 200 (x10) |

a.a.: all atoms; bb.: backbone

The resulting GaMD simulations produced a variety of transient binding modes, many resulting in partial dissociation of FASP. The high flexibility of FASP in these simulations suggests that our initial model lies in an unstable region of the binding free energy landscape (Figure S11A). This is not surprising considering our model was built based on a primed peptide (Tap63α) with a phosphate group (position -3) inserted in the first anion site, whereas in the FASP model this position is occupied by a positively charged lysine (see Figure S10B). In a significant portion of the trajectories, however, FASP rearranged into stably bound conformations and, in 14% of these, it adopted conformations prone to catalysis, with the oxygen of S662 near the γ phosphorous atom of ATP (Figure S11B). During the GaMD simulations, the activation loop remained mostly in the 'down' conformation and was particularly stable in the trajectories leading to bound states (replicas 1, 2 and 5 in Figure S11C).

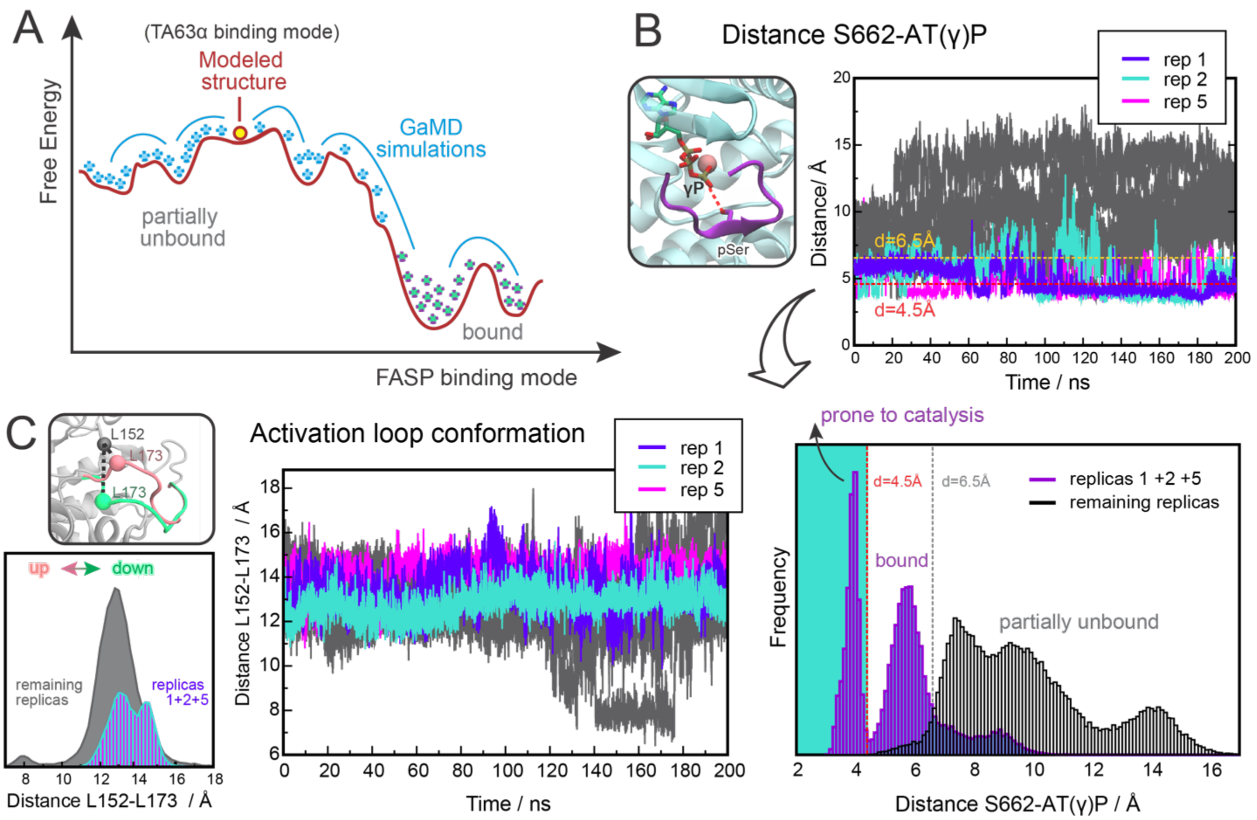


Figure S11. GaMD model of FASP-bound CK1. A) Schematic representation of FASP-CK1 binding landscape based on GaMD trajectories. B) Temporal evolution (top) and corresponding distribution (bottom) of the distance between S662 (γ oxygen) and the γ phosphorous atoms from ATP. C) Temporal evolution (right) and corresponding distribution (left) of the distance between γ-carbon atom of $Leu^{173}$ and the α-carbon of $Leu^{152}$. Trajectories that produced bound states (replicas 1, 2, and 5) are highlighted.

#### Final refinement with conventional MD simulations

To refine the binding mode of FASP, we selected 10 equally spaced conformations from trajectories that produced stably bound states as starting points for cMD simulations (Figure S12A). Based on these starting points, we ran 50 independent replicas (40 ns each), totaling 2µs of simulation time (Figure S12B). All replicas started with different initial velocities and used the same force field and parameters as described for the GaMD simulations, but without any boosting potential. Although FASP retained mobility freedom in the cMD simulations, it remained bound to CK1 and often sampled conformations that are prone to catalysis (Figure S12C). A detailed look at FASP dynamics during individual replicas (Figure S12D) reveals that FASP can rearrange in and out of its 'catalytically prone' binding mode within the nanosecond timescale.

Visual inspection of frames belonging to the 'catalytically prone' state reveals that the C-terminal portion of the FASP peptide retains high mobility freedom and does not engage in specific interactions with CK1. V663 at position +1 (relative to the priming serine) remains stably bound in the hydrophobic pocket between helix αF ($Tyr^{225}$) and the activation loop ($Leu^{173}$) (Figure S13A). The first anion binding pocket is partially occupied by E661 at position -1, which engages in electrostatic interactions with one of the positively charged clamps forming this site ($Arg^{178}$, in the P+1 loop) (Figure S13B). After moving out of the first anion site, K659 at position -3 engages in electrostatic interactions with the β phosphate of ATP and, at times, with Asp[132] (Figure S13C). Observations for the catalytically prone conformations were confirmed by RMSF and distance analyses (Table S6) reported in Figure 4 of the main manuscript.

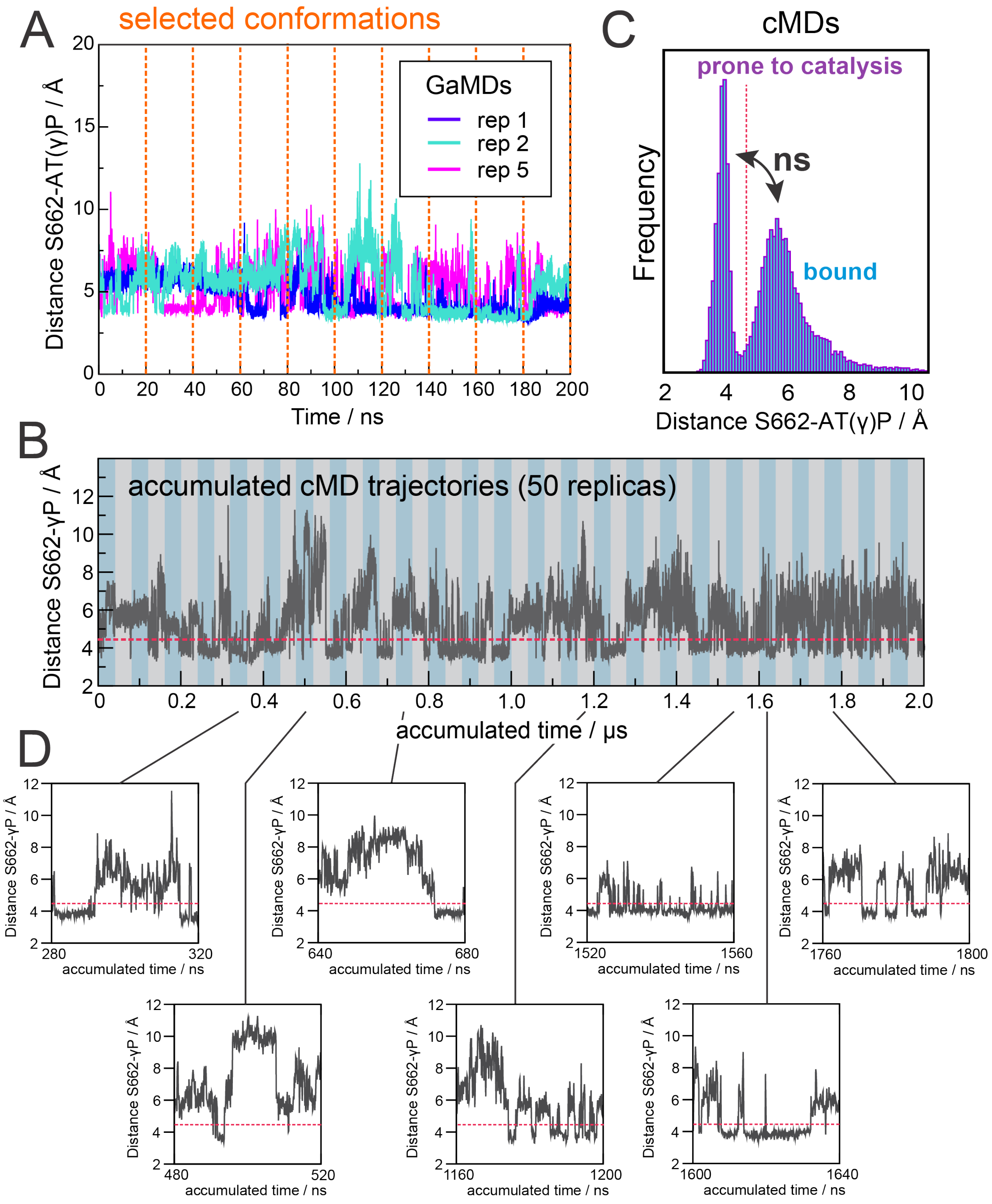


Figure S12. Final (cMD refined) model of FASP-bound CK1. A) Snapshots extracted from GaMD trajectories to be used as starting points to run cMD simulations. B) Distance between S662 (γ oxygen) and the γ phosphorous atoms from ATP calculated during the cMD simulations (each shaded stripe represents an individual replica). C) Distribution of the distances between S662 and AT(γ)P in the accumulated cMD simulations. D) Examples of binding mode transitions within individual cMD replicas.

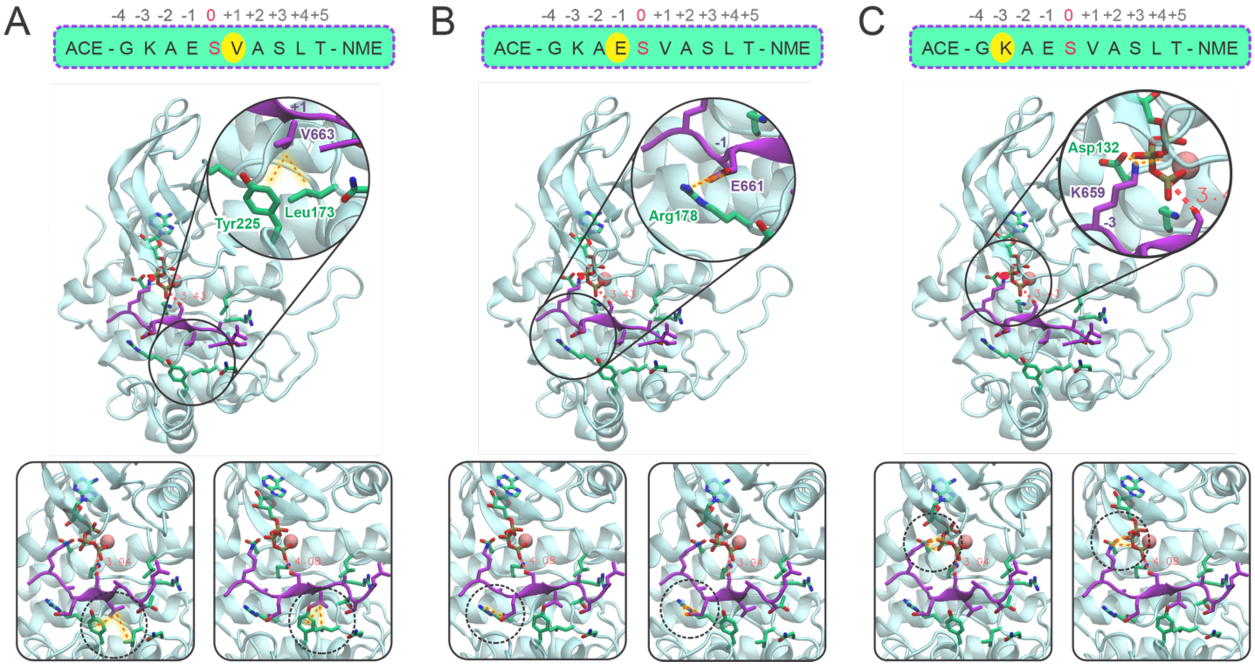


Figure S13. MD snapshot illustrating the interactions displayed between CK1 and FASP. A - C) Electrostatic interactions between V663 at position +1 (A), E661 at position -1 (B), and K659 at position -3 (C) of FASP with CK1.

Table S6. Definitions of how the distances reported in Figure 4 of the main manuscript were calculated.

| Distance | Selection 1 | Selection 2 |
|---|---|---|
| V663 to $Tyr^{225}$ | Res 663 (FASP) γ-carbons | Res 225 (CK1) γ-carbon, δ-carbons and ε-carbons |
| V663 to $Leu^{173}$ | Res 663 (FASP) γ-carbons | Res 173 (CK1) δ-carbons |
| E661 to $Arg^{178}$ | Res 661 (FASP) ε-oxygens | Res 178 (CK1) η-nitrogens |
| E661 to $Lys^{224}$ | Res 661 (FASP) ε-oxygens | Res 224 (CK1) ζ-nitrogen |
| K659 to AT(β)P | Res 659 (FASP) ζ-nitrogen | ATP and γ-phosphate |
| K659 to $Asp^{132}$ | Res 659 (FASP) ζ-nitrogen | Res 132 (CK1) δ-oxygens |

#### D. Biochemical validation of CK1-FASP model with mutational experiments

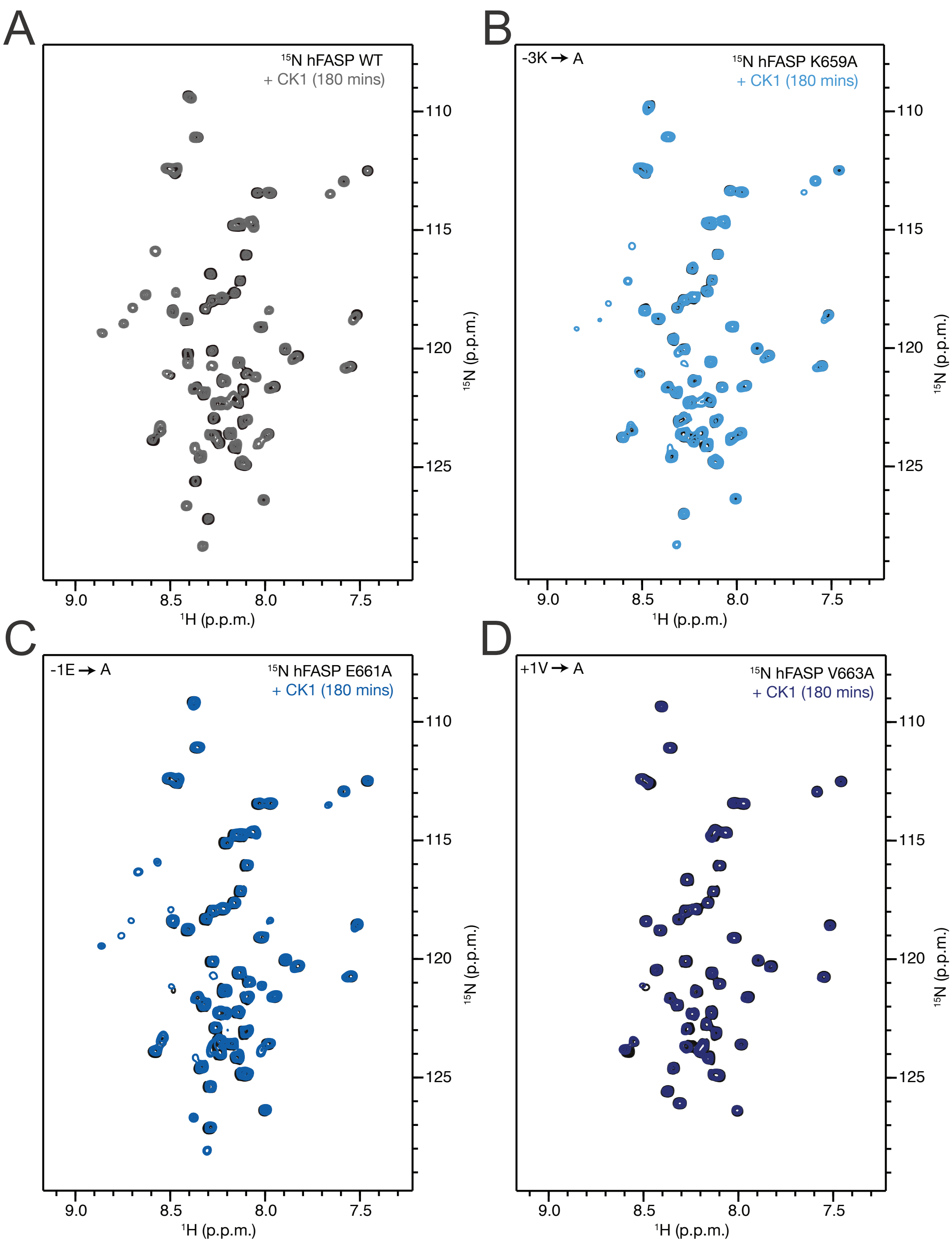


Figure S14. Biochemical validation of residues necessary for FASP priming. A - D) $^{15}N$-$^{1}H$ HSQC spectra comparing CK1 activity on human PER2 FASP WT (A) and alanine mutant FASP peptides K659A (B), E661A (C), and V663A (D) at a 3 hour timepoint.

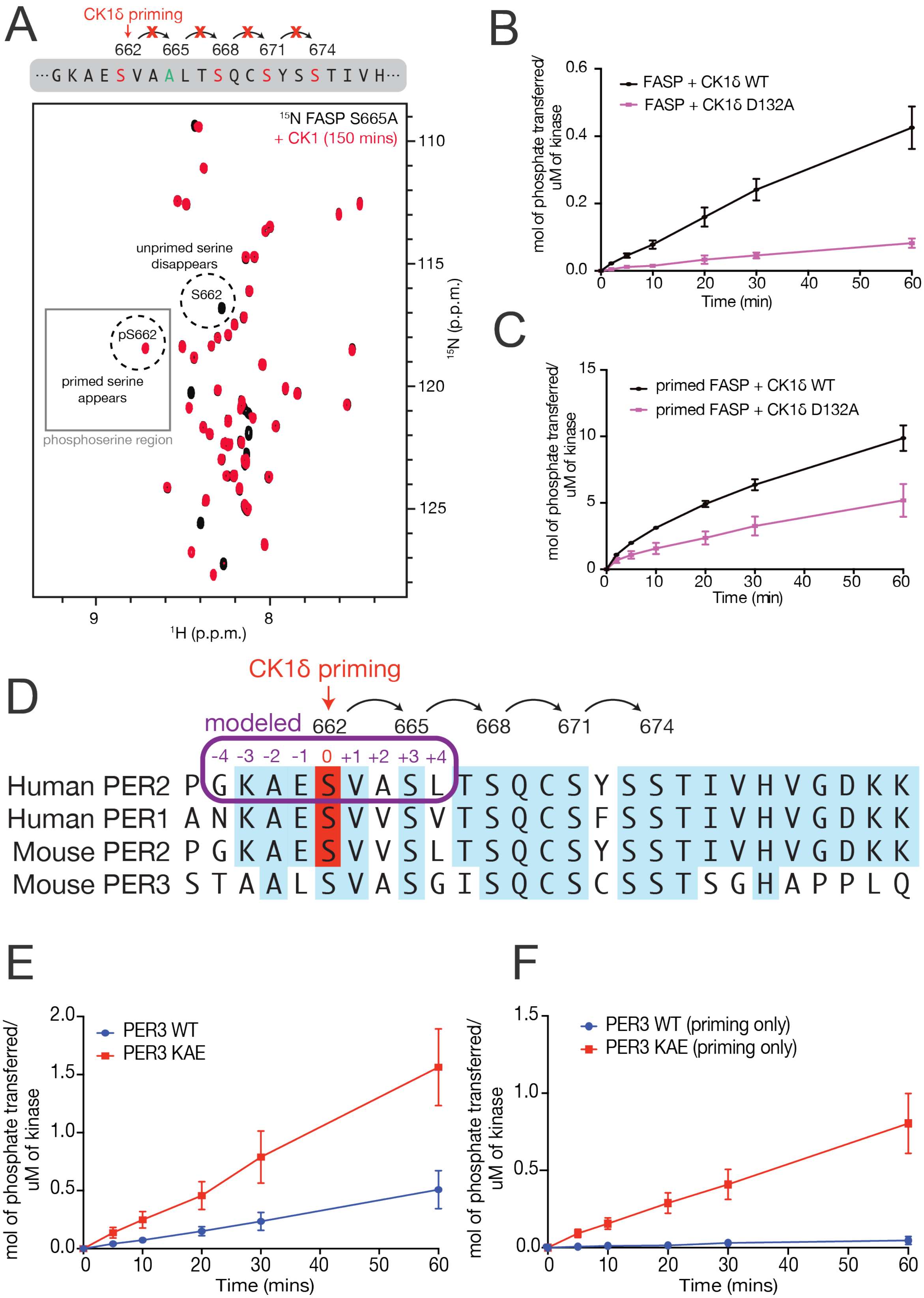
A
CK1δ priming
662 665 668 671 674
···G K A E S V A A L T S Q C S Y S S T I V H···
¹⁵N FASP S665A
+ CK1 (150 mins)
unprimed serine disappears
S662
pS662
primed serine appears
phosphoserine region
110
115
120
125
¹⁵N (p.p.m.)
9
8
¹H (p.p.m.)
B
mol of phosphate transferred/ uM of kinase
FASP + CK1δ WT
FASP + CK1δ D132A
Time (min)
C
mol of phosphate transferred/ uM of kinase
primed FASP + CK1δ WT
primed FASP + CK1δ D132A
Time (min)
D
CK1δ priming
modeled
662 665 668 671 674
-4 -3 -2 -1 0 +1 +2 +3 +4
Human PER2 P G K A E S V A S L T S Q C S Y S S T I V H V G D K K
Human PER1 A N K A E S V V S V T S Q C S F S S T I V H V G D K K
Mouse PER2 P G K A E S V V S L T S Q C S Y S S T I V H V G D K K
Mouse PER3 S T A A L S V A S G I S Q C S C S S T S G H A P P L Q
E
mol of phosphate transferred/ uM of kinase
PER3 WT
PER3 KAE
Time (mins)
F
mol of phosphate transferred/ uM of kinase
PER3 WT (priming only)
PER3 KAE (priming only)
Time (mins)

Figure S15. Biochemical validation of FASP priming model. A) Schematic depicting disruption of sequential kinase activity in the "priming only" human PER2 FASP peptide (S665A), along with $^{15}N$-$^{1}H$ HSQC spectra comparing activity between the WT and the "priming only" peptides at the 150 min timepoint of a CK1 kinase reaction showing no detectable downstream phosphorylation activity within the S665A peptide. B-C) $^{32}P$-ATP timecourse kinase reactions with synthetic mouse PER2 FASP peptides comparing activity on unprimed (B) and primed (C) substrates, between CK1 WT (black) and D132A (purple). D) Multiple sequence alignment of the FASP region from mammalian PER homologs with residue numbers corresponding to the human PER2 sequence. The -4 to +4 positions are labeled with respect to the priming serine (S662, red) and the purple box highlights the model peptide sequence from MD simulations. Blue shading indicates conservation of 75% or higher. E-F) $^{32}P$-ATP timecourse kinase reactions with synthetic mouse FASP peptides corresponding to the WT PER3 sequence (blue) and a PER3 sequence with the -3 and -1 residues mutated to the corresponding PER2 residues at those positions (-3K and -1E, respectively)(red). F) $^{32}P$-ATP timecourse kinase reaction as in panel E with serine residues downstream of the priming site mutated to alanine.

#### E. Reshaping of the substrate binding cleft and identification of allosteric binding pockets on the CK1 surface

To map binding sites on the surface of CK1, we employed FTMap [25], a fast computational approach that uses small organic probes to identify consensus sites (or pockets) that are likely to bind drug-like molecules. We submitted a series of conformations representative of the most populated MSM states to the FTMap server (https://ftmap.bu.edu/) and analyzed the results with FTProd [26]. For the WT system, we submitted 6 conformations from state I (loop up) and 21 conformations from state III (loop down). For the *tau* mutant, we submitted 26 conformations from state I' (loop up) and 21 conformations from state IV' (loop down with loop EF unfolded). Tables S7 and S8 describe the consensus sites (CS) detected for WT and *tau* conformations, respectively. When a site is detected (colorful cells), it contains a score based on the number of organic probes that docked in that site. When a cell is left blank, it means that FTMap did not detect that pocket in the corresponding CK1 structure.

Table S7. Consensus sites (CS) detected for WT CK1.

| State | # | CS0 | CS1 | CS2 | CS3 | CS4 | CS5 | CS6 | CS7 | CS8 | CS9 | CS10 |
|---|---|---|---|---|---|---|---|---|---|---|---|---|
| I (up) | 1 | 38 | - | 3 | - | 19 | - | 7 | - | 4 | 3 | 4 |
| I (up) | 2 | 32 | 5 | 2 | - | 21 | - | 15 | - | - | 7 | - |
| I (up) | 3 | 51 | - | 2 | 10 | 4 | - | 11 | - | 2 | 4 | - |
| I (up) | 4 | 44 | 8 | - | 7 | 12 | 2 | - | - | 4 | 6 | - |
| I (up) | 5 | 58 | 3 | 2 | - | - | - | - | - | 7 | 12 | - |
| I (up) | 6 | 45 | - | 5 | - | 6 | - | 7 | - | 4 | 16 | - |
| III (down) | 1 | 60 | - | - | - | - | 3 | - | - | 3 | 15 | - |
| III (down) | 2 | 36 | - | 23 | - | 4 | 9 | 3 | - | - | - | 2 |
| III (down) | 3 | 33 | 41 | 3 | 7 | - | - | - | - | - | - | - |
| III (down) | 4 | 42 | 19 | 5 | - | - | 5 | 2 | - | - | 6 | - |
| III (down) | 5 | 18 | 27 | 26 | - | 4 | - | 8 | - | - | - | - |
| III (down) | 6 | 14 | 21 | 20 | 4 | 21 | - | 4 | - | - | 2 | - |
| III (down) | 7 | 18 | 49 | - | 2 | 3 | - | 7 | - | - | 2 | - |
| III (down) | 8 | 39 | 10 | - | - | 3 | 13 | 6 | - | - | 11 | 3 |
| III (down) | 9 | 20 | - | 17 | - | 17 | - | 11 | - | 12 | - | - |
| III (down) | 10 | 47 | 25 | 9 | - | - | - | - | - | - | - | - |
| III (down) | 11 | 21 | 38 | - | 8 | - | - | - | - | - | 14 | 2 |
| III (down) | 12 | 22 | 38 | 18 | - | - | - | 5 | - | - | - | - |
| III (down) | 13 | 22 | 43 | 12 | - | - | - | - | - | - | 4 | - |
| III (down) | 14 | 7 | 33 | 24 | 5 | - | - | 2 | - | 6 | 3 | - |
| III (down) | 15 | 15 | - | 24 | 33 | - | - | - | - | - | 3 | - |
| III (down) | 16 | 18 | 26 | 14 | - | 6 | - | - | - | 14 | 3 | - |
| III (down) | 17 | 8 | 11 | 33 | 14 | 3 | - | 2 | 3 | 2 | 2 | - |
| III (down) | 18 | 34 | 18 | 17 | - | 3 | - | 7 | - | - | 2 | - |
| III (down) | 19 | 34 | 18 | 19 | - | 2 | 3 | - | - | 3 | - | - |
| III (down) | 20 | 38 | 3 | 30 | - | 5 | - | 6 | - | - | - | - |
| III (down) | 21 | 27 | 42 | 4 | - | 3 | - | 2 | - | - | - | - |

Table S8. Consensus sites (CS) detected for *tau* CK1.

| State | # | CS0 | CS1 | CS2 | CS3 | CS4 | CS5 | CS6 | CS7 | CS8 | CS9 | CS10 | C11 |
|---|---|---|---|---|---|---|---|---|---|---|---|---|---|
| III’ (up) | 1 | 33 | 23 | - | - | - | 7 | - | 3 | - | 4 | - | 6 |
| III’ (up) | 2 | 28 | 30 | - | 11 | 8 | - | - | - | - | - | - | - |
| III’ (up) | 3 | 20 | 31 | - | 2 | 16 | - | - | - | - | 12 | - | - |
| III’ (up) | 4 | 34 | 25 | - | 2 | 10 | 9 | - | - | - | - | - | 2 |
| III’ (up) | 5 | - | 23 | - | 21 | - | 24 | - | 3 | 4 | 5 | 2 | - |
| III’ (up) | 6 | 4 | 40 | - | 3 | 24 | 7 | - | - | - | 5 | - | - |
| III’ (up) | 7 | 13 | 27 | - | 31 | - | 10 | - | - | - | - | - | - |
| III’ (up) | 8 | 28 | 20 | - | 4 | 27 | - | - | - | - | - | - | - |
| III’ (up) | 9 | 15 | 38 | 15 | 3 | 6 | - | - | - | - | - | 3 | - |
| III’ (up) | 10 | 16 | 22 | - | 27 | - | 12 | - | - | - | - | - | 2 |
| III’ (up) | 11 | 36 | 28 | - | 4 | 8 | - | - | - | - | - | - | - |
| III’ (up) | 12 | 17 | 24 | 5 | 2 | 16 | - | - | 10 | - | 2 | - | - |
| III’ (up) | 13 | 17 | 24 | 5 | 2 | 16 | - | - | 10 | - | 2 | - | - |
| III’ (up) | 14 | 6 | 32 | - | - | 15 | 25 | - | - | - | 4 | - | - |
| III’ (up) | 15 | 37 | 26 | - | 13 | - | 3 | - | - | - | - | - | - |
| III’ (up) | 16 | 32 | 36 | - | 5 | 8 | - | - | - | 2 | 4 | - | - |
| III’ (up) | 17 | 19 | 16 | 7 | - | 17 | 5 | - | - | - | 16 | - | - |
| III’ (up) | 18 | 29 | 15 | 4 | 4 | 15 | 3 | - | - | 6 | - | 5 | - |
| III’ (up) | 19 | 33 | 26 | 2 | - | - | - | - | 8 | - | 9 | 2 | - |
| III’ (up) | 20 | 9 | 21 | - | 10 | 31 | 2 | - | - | - | 5 | - | - |
| III’ (up) | 21 | 37 | 39 | - | 4 | - | - | - | - | - | - | - | - |
| III’ (up) | 22 | 10 | 16 | - | 26 | 11 | - | - | - | 8 | 14 | - | - |
| III’ (up) | 23 | 45 | 17 | 8 | 4 | 4 | - | - | - | - | - | - | - |
| III’ (up) | 24 | 32 | 23 | - | 17 | 8 | - | - | - | - | - | - | - |
| III’ (up) | 25 | - | 24 | 16 | 19 | 12 | 5 | - | - | - | - | - | - |
| III’ (up) | 26 | 45 | 21 | - | - | 5 | 9 | 2 | - | - | - | - | - |
| IV’ (down) | 1 | 45 | 17 | 8 | 4 | 4 | - | - | - | - | - | - | - |
| IV’ (down) | 2 | 17 | 17 | - | 11 | - | 20 | 7 | - | - | 4 | - | - |
| IV’ (down) | 3 | 21 | 41 | 2 | 15 | - | - | - | - | - | - | - | - |
| IV’ (down) | 4 | 28 | 12 | - | 28 | 7 | 4 | - | - | - | - | - | - |
| IV’ (down) | 5 | 21 | 37 | 4 | 7 | 3 | 8 | - | - | - | 2 | - | - |
| IV’ (down) | 6 | 18 | 18 | 2 | 11 | - | 29 | - | - | - | 2 | - | 2 |
| IV’ (down) | 7 | 39 | 16 | 7 | - | - | 4 | - | - | 4 | - | 6 | - |
| IV’ (down) | 8 | 43 | 19 | - | 12 | - | 7 | - | - | 2 | - | - | - |
| IV’ (down) | 9 | 33 | 22 | 5 | 15 | - | 4 | - | - | - | 2 | - | - |
| IV’ (down) | 10 | 15 | 21 | 7 | - | 4 | 27 | - | - | - | 5 | - | 3 |
| IV’ (down) | 11 | 18 | 26 | - | 22 | - | 3 | - | - | - | 15 | - | - |
| IV’ (down) | 12 | 29 | 23 | 7 | 4 | - | - | - | - | - | 13 | - | 5 |
| IV’ (down) | 13 | 40 | 15 | 5 | 10 | 7 | 7 | - | - | - | - | - | - |
| IV’ (down) | 14 | 41 | 22 | 5 | 7 | - | 2 | - | - | - | - | - | - |
| IV’ (down) | 15 | 35 | 24 | - | 6 | 6 | 9 | - | - | - | - | - | - |
| IV’ (down) | 16 | 23 | 19 | 4 | 5 | - | 17 | - | - | 9 | 3 | - | - |
| IV’ (down) | 17 | 27 | 24 | 3 | 21 | - | - | - | - | 3 | - | - | - |
| IV’ (down) | 18 | 27 | 22 | - | 11 | 8 | - | 8 | - | - | - | 8 | - |
| IV’ (down) | 19 | 24 | 5 | 18 | - | 11 | 8 | 3 | 4 | - | 2 | - | - |
| IV’ (down) | 20 | 43 | 12 | 2 | 2 | - | 6 | - | - | 6 | 8 | - | - |
| IV’ (down) | 21 | 43 | 13 | - | 10 | 6 | - | - | 5 | - | 3 | 2 | - |

Table S9 describes the composition and location of the pockets detected in WT CK1, also illustrated in Figure S16. Not surprisingly, the top-scored pockets recapitulate the substrate binding cleft + active site (CS0), the ATP binding site (CS1) and the $Mg^{+2}$ binding site (CS2), showing that FTMap is able to identify these functional pockets even in apo structures. Interestingly, the ATP and the $Mg^{2+}$ binding sites are more likely to be closed or disassembled when the activation loop is up in the WT CK1 (compare scores and frequency of occurrence in ‘up’ and ‘down’ states). The next pocket detected in WT CK1 (CS3) is formed at the bottom of the kinase, between helices aG and aH.

Table S9. Composition and location of consensus sites identified in WT CK1.

| CS | Description | Residues | $\langle score\rangle^{Up}$ | $\langle score\rangle_{Down}$ | $Occurrence^{Up}$ | $Occurrence_{Down}$ |
|---|---|---|---|---|---|---|
| 0 | Substrate binding cleft + active site | 16 17 18 19 20 21 22 23 38 48 52 88 90 93 98 125 126 127 128 129 130 131 132 133 134 147 148 149 150 151 152 153 154 171 172 173 174 175 176 177 178 179 180 181 184 185 190 194 195 198 205 206 213 214 216 217 221 222 224 225 228 229 | 44.7 | 27.3 | 100% | 100% |
| 1 | ATP binding site | 13 15 16 17 23 24 25 36 37 38 52 56 66 80 81 82 83 84 85 86 87 88 89 90 91 130 132 133 134 135 138 148 149 150 151 | 5.3 | 27.2 | 50% | 80% |
| 2 | $Mg^{2+}$ binding site | 16 17 18 19 20 21 22 23 38 39 40 44 45 46 47 48 49 52 55 56 130 132 148 149 150 151 152 153 | 2.8 | 18.1 | 83% | 80% |
| 3 | FASP binding pocket | 180 182 185 197 201 213 232 233 234 235 236 237 240 249 253 256 | 8.5 | 10.4 | 33% | 33% |
| 4 | Back of the N-lobe | 34 60 31 62 63 64 65 66 67 69 83 84 85 137 138 141 144 146 | 12.4 | 6.2 | 83% | 57% |
| 5 | C-terminal pocket | 99 100 101 102 105 204 206 207 208 210 211 212 240 242 243 244 245 248 | 2 | 6.6 | 17% | 24% |
| 6 | Hinge pocket | 13 84 85 86 87 92 136 137 138 139 142 143 286 287 288 291 | 10 | 5 | 67% | 62% |
| 7 | 2nd anion binding site | 125 165 171 189 190 | --- | 3 | --- | 5% |
| 8 | 'Activation' pocket | 48 51 52 55 58 122 123 124 127 150 151 152 153 154 155 | 4.2 | 6.7 | 83% | 29% |
| 9 | Back of C-lobe | 63 106 107 110 111 114 115 139 140 141 142 143 | 8 | 16 | 100% | 76% |
| 10 | Back of C-lobe | 63 65 114 115 118 270 | 4 | 2.3 | 17% | 14% |

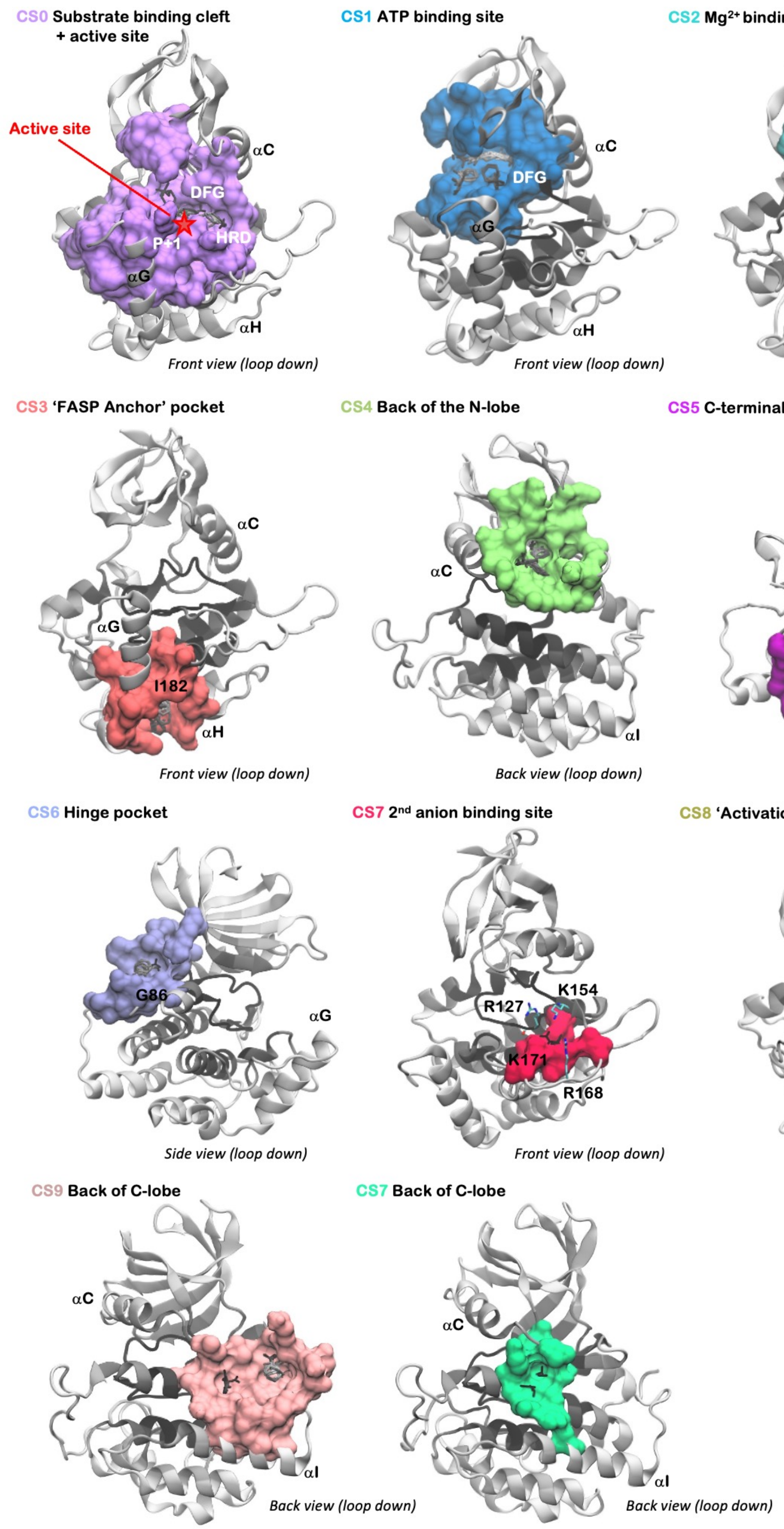


Figure S16. Consensus sites identified by FTMap on WT CK1.

Finally, Table S10 describes the composition and location of the pockets detected in *tau* CK1, also illustrated in Figure S17. As with WT CK1, the top scored pockets correspond to functional pockets including the ATP binding site (CS0) and the $Mg^{2+}$ binding site (CS1). However, we observe a fragmentation of the active site and substrate binding cleft, which are split into three pockets according to FTMap: the active site is absorbed into CS1 ($Mg^{2+}$ binding site), while the substrate binding cleft splits into CS3 (P+1 region) and CS5. The pocket located in the bottom between helices aG and aH is also detected and well scored in the *tau* mutant (CS2). Apart from these functional sites, the next pocket detected in *tau* (CS4) is located between the aC helix and the activation loop. It is noticeable that this pocket is more prominent when the activation loop is up (compare scores and frequency of occurrence in 'up' and 'down' states). An analogous pocket is also detected in the WT enzyme, but with poorer scores (see CS8 in Table S9), suggesting that this pocket is stabilized in the *tau* mutant.

Table S10. Composition and location of consensus sites identified in *tau* CK1.

| CS | Description | Residues | $\langle score\rangle^{Up}$ | $\langle score\rangle_{Down}$ | $Occurrence^{Up}$ | $Occurrence_{Down}$ |
|---|---|---|---|---|---|---|
| 0 | ATP binding site | 13 15 16 17 23 24 25 36 38 52 56 66 82 83 84 85 86 87 88 89 90 91 130 131 132 133 134 135 138 148 149 | 24.8 | 30.0 | 92% | 100% |
| 1 | $Mg^{2+}$ binding site + active site | 15 16 17 18 19 20 21 22 23 36 37 38 39 40 41 44 45 46 47 48 49 52 56 80 81 82 126 127 128 129 130 132 133 148 149 150 151 152 153 171 173 174 175 176 177 179 180 221 222 225 | 25.7 | 20.2 | 100% | 100% |
| 2 | FASP binding pocket | 182 197 201 204 213 232 233 234 235 236 237 240 252 253 256 | 7.8 | 5.6 | 31% | 67% |
| 3 | Substrate cleft (P+1 region) | 90 98 130 131 132 174 175 176 177 178 179 180 205 210 211 213 214 216 217 221 222 224 225 228 | 10.2 | 11.2 | 81% | 86% |
| 4 | 'Activation' pocket | 20 46 47 48 51 52 54 55 58 124 127 150 151 152 153 154 155 171 172 173 | 13.5 | 6.2 | 73% | 43% |
| 5 | Active site + part of the substrate cleft | 127 128 129 130 152 168 170 171 172 173 174 175 176 177 178 179 180 181 182 183 184 185 187 188 189 190 194 195 198 221 222 224 225 226 229 232 | 9.3 | 10.3 | 50% | 71% |
| 6 | 2nd anion binding site | 125 127 154 155 156 157 164 165 167 168 173 189 190 191 192 193 259 260 262 | 2.0 | 6.0 | 4% | 14% |
| 7 | Back of C-lobe | 61 62 63 64 65 66 67 83 114 115 117 118 141 146 | 6.8 | 4.5 | 19% | 10% |
| 8 | Back of N-lobe | 8 9 28 33 34 35 67 68 69 81 82 83 84 | 5.0 | 4.8 | 15% | 24% |
| 9 | Back of C-lobe | 106 107 108 109 110 114 115 140 141 142 143 144 270 273 274 277 278 284 286 288 | 6.8 | 5.4 | 46% | 52% |
| 10 | Hinge pocket | 86 87 92 136 137 139 140 142 143 290 291 | 3.0 | 5.3 | 15% | 14% |

| | | | | | | |
|---|---|---|---|---|---|---|
| 11 | C-terminal pocket | 93 98 99 100 102 105 206 207 244 | 3.3 | 3.3 | 12% | 14% |

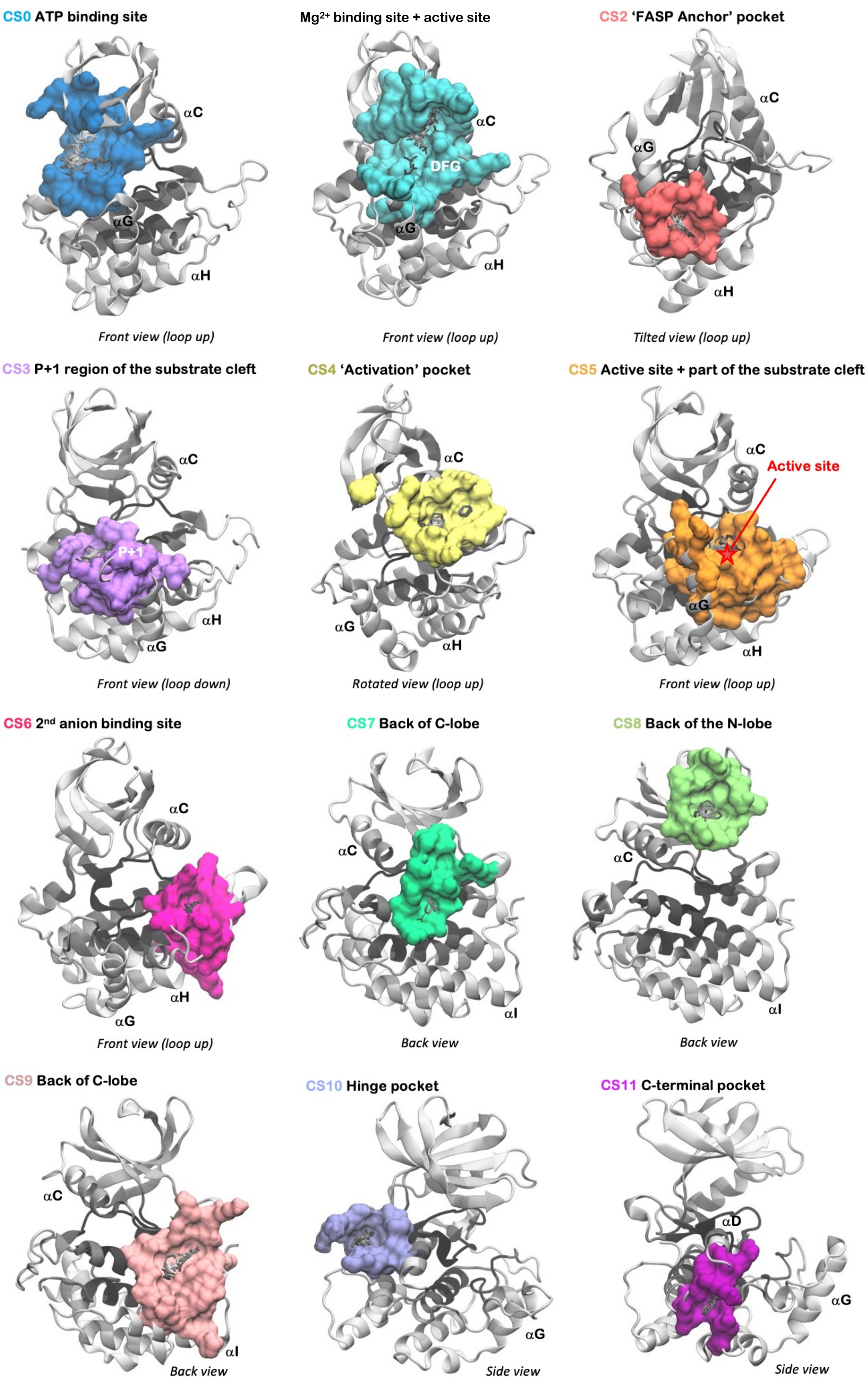


Figure S17. Consensus sites identified by FTMap on *tau* CK1.

#### References (for SI only)


1. Philpott, J.M., et al., *Casein kinase 1 dynamics underlie substrate selectivity and the PER2 circadian phosphoswitch.* Elife, 2020. **9**.
2. Jorgensen, W.L., et al., *Comparison of simple potential functions for simulating liquid water.* The Journal of Chemical Physics, 1983. **79**(2): p. 926-935.
3. Maier, J.A., et al., *ff14SB: Improving the Accuracy of Protein Side Chain and Backbone Parameters from ff99SB.* Journal of Chemical Theory and Computation, 2015. **11**(8): p. 3696-3713.
4. Wang, J., et al., *Development and testing of a general amber force field.* J Comput Chem, 2004. **25**(9): p. 1157-74.
5. Kashefolgheta, S. and A. Vila Verde, *Developing force fields when experimental data is sparse: AMBER/GAFF-compatible parameters for inorganic and alkyl oxoanions.* Phys Chem Chem Phys, 2017. **19**(31): p. 20593-20607.
6. Case, D.A., Betz, R.M., Cerutti, D.S., Cheatham III, T.E., Darden, T.A., Duke, R.E., Giese, T.J., Gohlke, H., Goetz, A.W., Homeyer, N., Izadi, S., Janowski, P., Kaus, J., Kovalenko, A., Lee, T.S., LeGrand, S., Li, P., Lin, C., Luchko, T., Luo, R., Madej, B., Mermelstein, D., Merz, K.M., Monard, G., Nguyen, H., Nguyen, H.T., Omelyan, I., Onufriev, A., Roe, D.R., Roitberg, A., Sagui, C., Simmerling, C.L., Botello-Smith, W.M., Swails, J., Walker, R.C., Wang, J., Wolf, R.M., Wu, X., Xiao, L., Kollman, P.A., *AMBER 2016*. 2016: University of California, San Francisco.
7. Darden, T., D. York, and L. Pedersen, *Particle mesh Ewald: An <i>N</i>·log(<i>N</i>) method for Ewald sums in large systems.* The Journal of Chemical Physics, 1993. **98**(12): p. 10089-10092.
8. Pérez-Hernández, G., et al., *Identification of slow molecular order parameters for Markov model construction.* Journal of Chemical Physics, 2013. **139**(1).
9. Scherer, M.K., et al., *PyEMMA 2: A Software Package for Estimation, Validation, and Analysis of Markov Models.* Journal of Chemical Theory and Computation, 2015. **11**(11): p. 5525-5542.
10. Wehmeyer, C., et al., *Introduction to Markov state modeling with the PyEMMA software [Article v1.0].* Living Journal of Computational Molecular Science, 2019. **1**(1): p. 5965.
11. Prinz, J.H., J.D. Chodera, and F. Noé, *Spectral Rate Theory for Two-State Kinetics.* Physical Review X, 2014. **4**(1).
12. Röblitz, S. and M. Weber, *Fuzzy spectral clustering by PCCA plus : application to Markov state models and data classification.* Advances in Data Analysis and Classification, 2013. **7**(2): p. 147-179.
13. Deuflhard, P. and M. Weber, *Robust Perron cluster analysis in conformation dynamics.* Linear Algebra and Its Applications, 2005. **398**: p. 161-184.
14. Kube, S. and M. Weber, *A coarse graining method for the identification of transition rates between molecular conformations.* Journal of Chemical Physics, 2007. **126**(2).
15. Noé, F., et al., *Projected and hidden Markov models for calculating kinetics and metastable states of complex molecules.* Journal of Chemical Physics, 2013. **139**(18).
16. Chodera, J.D., et al., *Bayesian hidden Markov model analysis of single-molecule force spectroscopy: Characterizing kinetics under measurement uncertainty.* arXiv preprint arXiv:1108.1430, 2011.
17. Ho, B.K. and R. Brasseur, *The Ramachandran plots of glycine and pre-proline.* Bmc Structural Biology, 2005. **5**.
18. Gebel, J., et al., *p63 uses a switch-like mechanism to set the threshold for induction of apoptosis.* Nat Chem Biol, 2020. **16**(10): p. 1078-1086.
19. Martínez, L., R. Andreani, and J.M. Martínez, *Convergent algorithms for protein structural alignment.* Bmc Bioinformatics, 2007. **8**.
